# Sensory and Motor Building Blocks of an Extra Robotic Digit Representation

**DOI:** 10.64898/2026.11.09.765353

**Authors:** Lucy Dowdall, Elena Amoruso, Giulia Dominijanni, Maria Molina-Sanchez, Jörn Diedrichsen, Dani Clode, Tamar R. Makin

**Affiliations:** MRC Cognition and Brain Sciences Unit, University of Cambridge, Cambridge, UK; Institute of Cognitive Neuroscience, University College London, London, UK; Neuro-X Institute, Ecole Polytechnique Fédérale de Lausanne, Lausanne, Switzerland; Western Institute of Neuroscience, Western University, London, Ontario, Canada; Department of Computer Science, Western University, London, Ontario, Canada; Department of Statistical and Actuarial Sciences, Western University, London, Ontario, Canada; Dani Clode Design, Cambridge, UK

## Abstract

Supernumerary limbs provide a unique opportunity to ask how the brain represents an entirely novel effector within an existing sensorimotor system. Here, we investigated how primary motor cortex (M1) represents a robotic extra digit (the Third Thumb; Dani Clode Design), and how this representation changes with experience. Using fMRI, we compared participants following short-term exposure with participants who had undergone longitudinal Third Thumb training. The device was controlled by the toes but operated alongside the biological fingers, while its sensory consequences were experienced on the hand. Following prolonged experience, greater activity was elicited in the M1 hand area both during Third Thumb control, and in response to its sensory consequences alone. Multivariate analyses showed that the Third Thumb representation became more distinct from its biological toe controller. Pattern component modelling further showed that the representation could be explained by a combination of its constituent motor and sensory patterns, but their relative contributions changed substantially with experience, from toe-dominated following short-term exposure to sensory-dominated following longitudinal training. These findings suggest that M1 can represent a novel effector through flexible recombination of existing sensorimotor representations, with experience altering how these pre-existing resources contribute to the emerging representation.

## Introduction

The brain contains a highly detailed body map. In primary sensorimotor cortex, neighbouring body parts are broadly arranged according to a somatotopic map. Within this organisation, individual effectors are represented by distributed and partly overlapping patterns of activity associated with their movement and sensation (Dall’Orso et al., 2018; Muret et al., 2022).

The plasticity of this cortical map has long been the subject of study. Our sensorimotor body representation is established from early in development (Dall’Orso et al., 2018; Meltzoff et al., 2019), and remarkably stable (Arbuckle et al., 2019; Kolasinski et al., 2016): It is maintained in the case of lost sensorimotor function (Howell et al., 2026; Krakauer & Carmichael, 2017; Mancini et al., 2019) and even amputation (Kikkert et al., 2016; Schone et al., 2025; Wesselink et al., 2019). Despite this large-scale stability, there are also many reported examples of use-dependent plasticity in body representation (Ejaz et al., 2015; Kolasinski et al., 2016; Raffin & Siebner, 2019). This coexistence of large-scale stability and use-dependent changes therefore prompts a key question: just how malleable is our body map?

A key limitation when investigating such plasticity mechanisms, is that from birth we have a set bodily architecture. Consequently, we are restricted in the methods we can use to probe the body representation - we can only study the refinement of how we use our body (through sensorimotor skill acquisition; Ogawa et al., 2019; Raffin & Siebner, 2019; Wiestler & Diedrichsen, 2013), or its response to alterations such as sensory deprivation (Kikkert et al., 2021; Makin et al., 2013; Wesselink et al., 2022). Even in individuals with a congenital limb difference (Hahamy et al., 2017; Maimon-Mor et al., 2021; Wesselink et al., 2019), the body topography is set from an early age (Tucciarelli et al., 2026), providing limited further insight in the plasticity of the body representation through life experience.

Augmentation technology, in particular supernumerary limbs, therefore provide a unique opportunity. These technologies are designed to extend and enhance the functionality of an intact body (Eden et al., 2022; Prattichizzo et al., 2021; Wu & Asada, 2014). By (temporarily) introducing a novel effector, without replacing any existing body part, these technologies require users to learn entirely new ways of recruiting and coordinating their body. For example, following training by healthy participants to use an extra robotic digit (the ‘Third Thumb’; Dani Clode Design; Figure 1A), the natural finger synergies were updated as the device was integrated into their motor repertoire (Dowdall et al., 2026; Kieliba et al., 2021). These kinematic changes were accompanied by use-dependent changes to the representation of the biological fingers in primary motor cortex (M1) following Third Thumb training (Kieliba et al., 2021).

**Figure 1:**
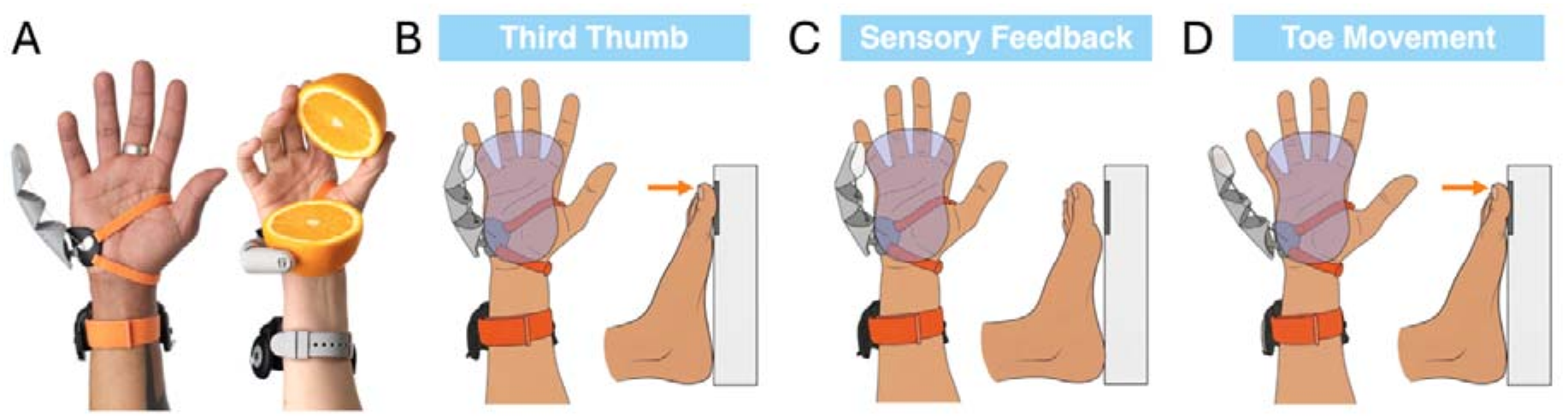
Third Thumb MRI task setup and primary experimental conditions. (A) The Third Thumb (Dani Clode Design) is a 3D-printed supernumerary robotic digit worn on the ulnar side of the hand. In its standard configuration, movement is proportionally controlled by force sensors positioned beneath the big toes, providing two degrees of freedom. (B-D) Main experimental conditions used to dissociate the sensory and motor components of Third Thumb control. (B) During Third Thumb movement, participants pressed the right big toe to move the Third Thumb against the object, producing corresponding intrinsic somatosensory feedback on the hand. (C) During passive sensory feedback, the Third Thumb moved automatically against the object while participants remained still, providing the sensory consequences of Third Thumb movement without active toe control. (D) During toe movement, participants pressed the right big toe against the footrest while the Third Thumb remained inactive, isolating movement of the biological controller.

But what about the device itself? How does the brain represent a novel supernumerary limb? The challenge here is that the sensorimotor system can only access the technology through pre-existing bodily channels; motor commands must be derived from existing control signals, while somatosensory feedback must be conveyed through established sensory pathways (Dominijanni et al., 2021). In the case of the Third Thumb (Clode et al., 2024; Kieliba et al., 2021), the device is controlled through movements of the toes, whereas the sensory consequences of its actions are collected on the hand. Because the Third Thumb is worn on the palm, forces arising from its contact with the environment are transmitted directly to the skin of the palm, naturally providing intrinsic somatosensory feedback about the device’s actions.

We previously found these haptic signals arising from the Third Thumb are immediately represented in primary somatosensory cortex (S1) in a topographically organised manner, which can then be refined through experience (Dowdall et al., 2026). However, because these signals are conveyed through the device’s attachment to the palm, this refinement reflects changes in how Third Thumb-related input is integrated within the existing hand representation, rather than the emergence of a dedicated representation of the device. These findings therefore leave unresolved whether the brain can construct a novel representation of new effector that is distinguishable from the biological channels through which it is controlled and sensed.

The Third Thumb provides a framework to examine this by providing a dissociation between motor control and function. By using motor cortex as scaffolding, we can ask whether the motor representation of the supernumerary limb is organised around the body part providing the control signal, the hand-related function it serves and where the associated sensory feedback is received, or whether a representation develops that is distinguishable from both. Successful use is possible from the first exposure (Clode et al., 2024), suggesting that existing sensorimotor resources can be rapidly co-opted to support device control. Following longitudinal training, performance continues to improve (Kieliba et al., 2021), and, importantly, this learning can generalise beyond the original sensorimotor mapping to a different hand and control signal (Molina-Sanchez et al., 2026). This generalisation suggests that learning is not restricted to the specific biological effectors used during training. Building on this and our previous research on expert prosthetic limb (Maimon-Mor & Makin, 2020) and tool-use (Schone et al., 2021) representation, we therefore predicted that the motor representation of the Third Thumb would initially be anchored to the body parts supporting its control but would become increasingly distinct from these components following extended experience.

In the present study, we assessed how the sensory and motor components may combine to construct the Third Thumb representation following short-term (up to 90 minutes) training and long-term (minimum of seven days) experience. While wearing and actively controlling an MRI-compatible Third Thumb in the scanner, participants performed a movement task that allowed identification of the specific toe-controller and sensory-feedback activity patterns to investigate how this information combines in primary motor cortex (M1) to represent the Third Thumb. To isolate differences associated with Third Thumb experience from familiarity with the specific scanner task, we used a between groups design, such that both the short- and long-term experience groups encountered the fMRI paradigm with comparable task-specific novelty.

## Results

### Primary motor cortex retunes to Third Thumb inputs following training

To explore the responses of the brain to the Third Thumb and its sensory and motor components, participants performed a tapping task in the MRI scanner while actively controlling an MRI-compatible Third Thumb. In separate conditions (Figure 1B-D), participants would:

i. Control the Third Thumb to tap on an object, placed on their hand. This involved pressing their right toe against a foot pedal, while also experiencing somatosensory feedback on their hand, resulting from the Thumb’s movement and subsequent impact when interacting with the object.
ii. Move their right toe (with no Thumb movement) to press against the foot pedal, to replicate the toe control (no Thumb movement).
iii. Passively experience the hand-worn Third Thumb moving autonomously to tap on the object (no toe movement). There were three additional control conditions involving the biological fingers:
iv. Move the biological thumb (D1) to tap on the object (no toe movement)
v. Move the biological thumb (D1) to tap on the object, while also moving the right toe to press against the foot pedal
vi. Move the biological little finger (D5) to tap on the object, while also moving the right toe to press against the foot pedal

The short-term and long-term training groups presented in this paper relied on two separate samples, acquired with the same paradigm in two different scanners. Our main analysis focused on group differences, see *Methods: Controlling for Between-Group Differences* for verification of comparable acquisition parameters, pre-processing, motion outliers and pattern reliability between groups.

Our primary interest was focused on motor control of the Third Thumb, motivating us to focus our analyses on the M1 hand area (Figure 2A; see Methods for details on region of interest (ROI)). We first considered M1 activity levels (Figure 2B) in response to each of our conditions following short-term experience. As expected, D1 movements activated the M1 hand area (activity significantly above zero; *t*(21)=9.962,*p*<.001), while toe movement did not significantly activate the region (*p*=.599,BF_10_=.254), nor did the passive Third Thumb movement (*p*=.160,BF_10_=.561). When actively moving the Third Thumb, there was significant deactivation of the M1 hand region (*p*=.035) compared to baseline, supporting that neural resources of the hand motor area are not recruited when using the Third Thumb after initial exposure.

**Figure 2:**
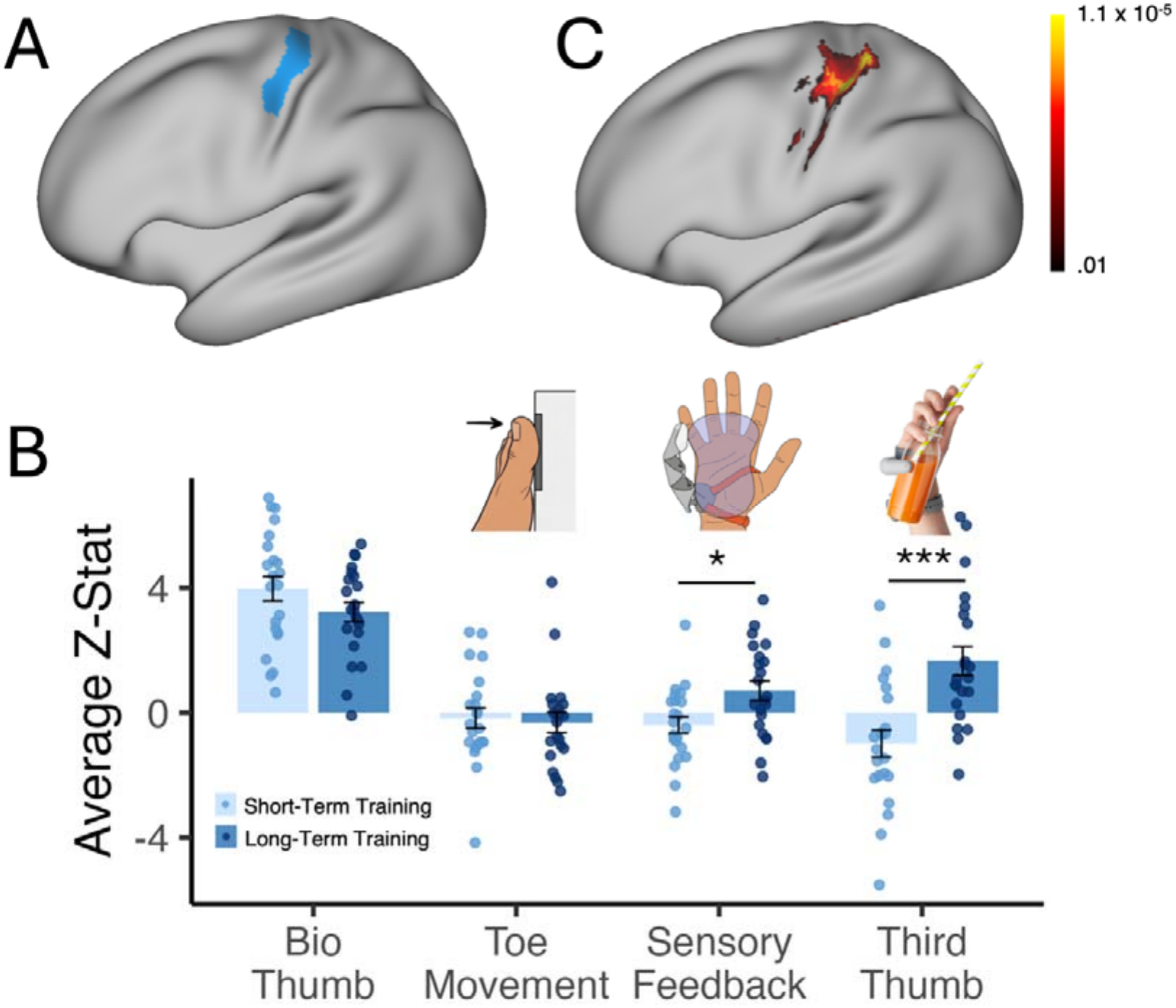
Experience-dependent recruitment of the M1 hand area during Third Thumb control. (A) M1 hand area region of interest (ROI) used for the primary analyses, centred around the anatomical hand knob (see Methods for full details of ROI definition). (B) Average activity within the M1 hand area ROI during biological thumb (D1) movement, toe movement, passive sensory feedback and active Third Thumb movement. Biological thumb movement produced strong M1 hand area activity in both groups, while toe movement did not differ between groups. Following long-term experience, activity was significantly greater during passive Third Thumb sensory feedback and during active Third Thumb movement, compared with the short-term group. Individual points represent participants. (C) Whole-brain contrast of activity during Third Thumb movement between the long- and short-term experience groups. Greater activity following long-term experience was centred around the sensorimotor hand knob; positive values indicate greater activity in the long-term relative to the short-term group, with lighter colours indicating higher p values. The map is displayed at p<.01, cluster-corrected at p<.05 * denotes p < .05; *** denotes p < .001.

We next examined whether this activity profile differs with experience in the second cohort of participants who had undergone longitudinal training with the Third Thumb. Moving D1 again produced the expected M1 activation (*t*(21)=10.30,*p*<.001), which was not significantly different from D1 activity found for the short-term training group (*t*(42)=1.472,*p*=.149,). The Bayes factor (BF_10_=.706) only implied anecdotal evidence for similar activation across the groups, such that we could not confidently conclude that the methodical differences between the groups did not play a role. Therefore, all further group comparisons were explored using the interaction terms relative to D1 (i.e. Group x Movement Condition interactions).

Similarly to the short-term experience group, there was no significant activity in the motor hand area for toe movements (*t*(21)=.990,*p*=.334,BF_10_=.345), with no significant group differences (Group x Condition interaction: *F*(1,42)=1.249,*p*=.270; group main effect: BF_10_ .310). However, following training, simply receiving the sensory feedback from passive Thumb movement now significantly activated the M1 hand area (t(21)=2.258,*p*=.035), with corresponding significantly higher activation following long-term experience (Group X Condition interaction *F*(1,42)=12.346,*p*=.001; short-vs long-term training, *p*=.019). Importantly, Third Thumb movement in the long-term training group resulted in positive activity in the hand motor area *t*(21)=3.587,*p*=.002, as reflected in highly significant group differences (*F*(1,42)=26.184,*p*<.001; short-vs long-term training, *p*<.001). Group differences were also evident when comparing this Third Thumb activity to the sum of the average toe and feedback activity (*F*(1,42)=10.294,*p*=.003). The summed activity was significantly below the Third Thumb activity level for the long-term group (*p*<.001) but not for the short-term group (*p*=.259,BF_10_ 1.079). Importantly, these experience-dependent effects were not observed in the M1 foot area (*p*=.129; see *Supplementary Materials*). Given that M1 foot area activity reflects the magnitude of toe-related information to the motor cortex, this suggests that the increased M1 hand area response is not simply a consequence of greater recruitment of the biological toe controller during Third Thumb movement. Together, these findings demonstrate an experience-dependent change in how the M1 hand area is recruited during Third Thumb-related sensorimotor processing.

To assess if there were any further experience-dependent differences during Third Thumb control that warranted exploration, a group whole-brain contrast between the longitudinal (minimum of seven days) and short-term experience (one session, 90 min) groups was conducted. This revealed a cluster of increased activity around the sensorimotor hand knob (Figure 2C). With this clear verification of our area of interest, we focused our resulting analyses on the Third Thumb representation in the motor hand area (see *Supplementary Materials* for complementary analyses in the primary somatosensory cortex (S1) hand area). The group whole-brain contrast additionally revealed clusters of activity in S1 and anterior IPS during the sensory feedback condition, as well as the lateral occipital cortex. No clusters of interest were found when contrasting the groups during toe movement (see Supplementary Materials S1 for group whole brain-contrast maps). Further exploratory analyses in premotor cortex did not produce any experience-dependent results of interest (see *Supplementary Materials* for univariate analyses in premotor dorsal and ventral).

### Experience-dependent differences in Third Thumb representational geometry

The univariate analysis informs us about differences in the recruitment of M1 hand area and the amplitude of the neural response when using the Third Thumb. To gain further insight into what is underlying these differences, we next used representation similarity analysis to compare the information content of the activity patterns elicited by each group. By looking at the patterns of the voxel-wise responses, this enables characterisation of the representational relationships between the Third Thumb and its sensory and motor components.

To quantify the organisation underlying the neural response to the Third Thumb condition, the similarity of its activity pattern was compared to both the other digits on the hand, and its respective components using cross-validated Mahalanobis (crossnobis) distances.

Before exploring the Third Thumb’s relationship to its components, we first established the stability of the Thumb representation in relation the biological hand. The Third Thumb representation was considered in relation to coordinated biological movements that matched either its function (coordinated D1 and Toe movement (D1&Toe)) or its anatomical position (coordinated D5 and Toe movement (D5&Toe)). Consistently across both groups, the distance between the Third Thumb and D5&Toe was larger than between the Third Thumb and D1&Toe *(F*(1,42)=58.26*,p*<.001; Figure S2). This suggests the Third Thumb is represented more similarly to its functional equivalent than its topographic neighbour. This was replicated when taking the distance between the passive sensory feedback and D1&Toe vs D5&Toe: *F*(1,42)=38.671,*p*<.001, where the feedback was again more similar to D1&Toe across groups. Between groups, the distance between D1 and the Third Thumb was not significantly different (*t*(42)=.316,*p*=.754,BF_10_=.319). This hints that any changes incurred to the Thumb representation through experience were not associated with the biological hand motor representation. Nevertheless, all further distances were still assessed in relation to the Third Thumb to D1 distance.

We next investigated the representational geometry between the Third Thumb and its two consistent toe movement and sensory feedback components, visualised in Figure 3A using a multidimensional scaling (MDS) plot in the M1 hand area. Based on the first two dimensions, there appears to be differences in the geometric relationship between the Third Thumb representation and its components following long-term (Figure 3A) experience compared to short-term (Figure 3B). In particular, the Third Thumb is becoming more dissimilar from the toe movement.

**Figure 3:**
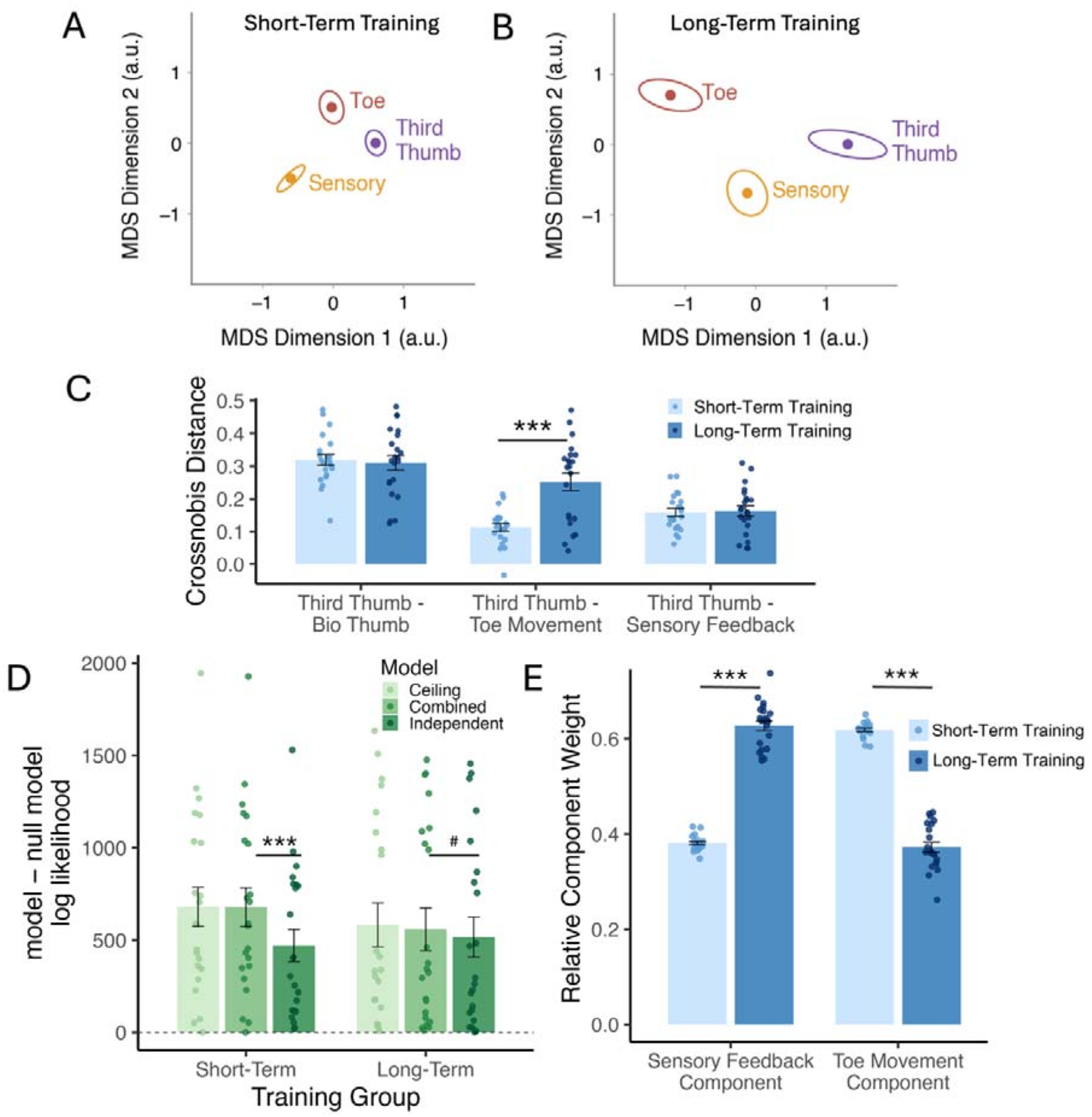
Representational geometry of the Third Thumb and its constituent components in the M1 hand area and their experience-dependent changes. Multidimensional scaling (MDS) visualisation of the representational relationships between Third Thumb movement, toe movement and passive sensory feedback following (A) short-term and (B) long-term Third Thumb experience. Relative positions were derived from the group-averaged cross-validated second-moment matrix (G) of the multivoxel activity patterns elicited by each condition, with greater separation indicating greater representational dissimilarity. Two-dimensional MDS projection is shown for visualisation only, configurations were rigidly reoriented for visual comparison, with the Third Thumb aligned to the positive x-axis and the toe representation positioned above it. (C) Representational similarity analysis within the M1 hand area. Cross-validated Mahalanobis (crossnobis) distances quantify the dissimilarity between the Third Thumb activity pattern and the biological thumb (D1), toe movement and passive sensory feedback patterns. The Third Thumb-toe distance was significantly greater following long-term experience. In contrast, the Third Thumb-biological thumb and Third Thumb-sensory feedback distances did not significantly differ between groups. (D) Pattern component modelling tested whether the Third Thumb activity pattern could be explained as a weighted combination of its constituent toe movement and sensory feedback patterns. Cross-validated log-likelihoods for each model (independent, combined and noise ceiling) are subtracted from a null model (for visualisation only), with higher values indicating better model fit. Following short-term experience, the combined model significantly outperformed the independent model, following long-term experience, evidence for this difference was inconclusive. The combined model did not differ significantly from the noise ceiling in either group, indicating that it provided an adequate account of the reliable Third Thumb representational structure. (E) Relative component weightings estimated from the combined model. Following short-term experience, Third Thumb-related activity was predominantly explained by the toe movement component. With long-term experience, this relationship reversed, with a significant increase in the sensory feedback weighting and corresponding decrease in the toe movement weighting. Individual dots represent participants. *** denotes p<.001, # denotes trending p-value (p=.056).

When assessing the distances between the Third Thumb and the toe movement and sensory feedback component following only short-term experience, the Third Thumb contained distinguishable information from both the toe movement (distance significantly above zero; *p*<.001) and the sensory feedback (*p*<.001). The distance between the Third Thumb and toe movement then became significantly greater following long-term experience compared to short (*F*(1,42)=34.18,*p*<.001; short vs long-term *t*(61.29)=4.865,*p*<.001; Figure 3C). The Third Thumb representation therefore became more dissociated from its biological motor component following experience, suggesting that in the M1 hand area, Third Thumb use becomes less dominated by its biological controller. Importantly, this increased separation from the toe condition was not accompanied by greater similarity to the passive Third Thumb movement (sensory feedback: *F*(1,42)=.325,*p*=.572; short vs long-term *t*(66.55)=.199,*p*=.843,BF_10_ .305; Figure 3C), hinting that experience may change how the available sensory and motor components are weighted and combined.

### Re-weighting of the sensory and motor components

To directly investigate how these constituent sensory and motor components may be re-combined with experience, pattern component modelling (PCM (Diedrichsen et al., 2018) was utilised. This modelled how the voxel-wise activity patterns for the toe movement and sensory feedback components may combine to predict the Third Thumb representation using four models – (i) combined model – assumes the Third Thumb representation can be explained by a weighted sum of the patterns associated with sensory feedback and toe movement (ii) independent model – assumes the Third Thumb representation is not related (orthogonal) to the two components (iii) noise ceiling model – a fully flexible model to assess quality of the fit from the mean of the other subjects (iv) null model – assumes there is no condition-relevant representational structure, used as a baseline for assessing model fits.

To assess model performance, the combined model was compared against the independent model (Figure 3D). The combined model performed better following short-term experience (*p*<.001; combined minus independent model log-likelihood *M*Δ*=*207.6 *(SD=*147.3)), whereas evidence for an advantage following long-term experience was inconclusive (*p*=.056,BF_10_=1.314; combined minus independent model log-likelihood *M*Δ*=*41.53 *(SD=*104.1)). With this borderline result, we then validated whether the combined model sufficiently explained the variance in the data for both groups by comparing its fit to the noise ceiling, of which it was not significantly different (short-term *p*=.449,BF_10_=.309; noise ceiling minus combined log-likelihood *M*Δ=1.802 (*SD*=11.35); long-term *p*=.203,BF_10_=.686; noise ceiling minus combined log-likelihood *M*Δ=24.92 (*SD*=77.31)). These model comparisons demonstrate the combined model provides an adequate account of the Third Thumb representation.

To understand how the constituent components combine to explain the Third Thumb representation, the relative weightings of each component in the model were assessed (Figure 3E). When comparing groups, we found a significant interaction (Group X Component; *F*(1,42)=534.447,*p*<.001). Following short-term experience, the toe movement was the dominant component explaining the Third Thumb representation (significantly higher weighting; *t*(21)=37,*p*<.001). However, following longitudinal experience, this pattern was reversed (*t*(21)=12.56,*p*<.001), resulting in a significant increase in the weighting of the sensory feedback component (*t*(41)=12.579,*p*<.001), and a corresponding significant decrease for the weighting of the toe movement (*t*(41)=12.579,*p*<.001). Therefore, in addition to the dissociation from the toe representation we observed above following experience, we also find increased utilisation of Third Thumb sensory input on the hand by the M1 hand area.

As a reference to consider how such sensorimotor components are normally weighted, a biological body combined model was structured that explained the coordinated movements of D1 and Toe with the components of D1 and Toe (see model fits and weights in Supplementary Figure S3). Overall model performance of this biological model was compared to the Third Thumb model, relative to the null. This revealed a significant interaction between group and condition (*F*(1,126)=27.011,*p*<.001). Follow-up contrasts demonstrated that while overall model performance was significantly higher for the biological than Third Thumb movement in both groups (both *p*<.001), this difference was smaller following long-term (EMM difference=765.3) compared to short-term (EMM difference=1459.6). Thus, the difference in performance for the Third Thumb and biological body models following short-term experience was somewhat reduced following prolonged training.

Finally, given the re-weighting of resources in the motor hand area, follow-up analyses added D1 to the model with the motor and sensory components to assess if incorporating this biological hand component improves the model fit. When comparing the original, two-component model to this extended model, a significant Group X Model interaction (*F*(1,42)=8.280,*p*=.006) was revealed. Including D1 significantly reduced model performance following short-term experience (*p*<.001), whereas no significant difference was observed following long-term training (*p*=.848, BF_10_≈.296). The model did not perform significantly better with the biological hand component, limiting any inference regarding relevance of D1 to the Third Thumb representation. However, this experience-dependent difference supports increased integration of the Third Thumb with biological hand resources following experience.

### Representational differences across the sensorimotor network

Lastly, the representational geometry of the Third Thumb and its components was explored across the whole brain to identify if there were further areas with differences in geometry following long-term training. Crossnobis distances between the Third Thumb and (i) toe movement, (ii) sensory feedback, and (iii) D1 were calculated within each of the cortical parcellations identified by the human connectome project (Glasser et al., 2016). The only distance pair that produced significant parcels of interest was the Third Thumb and toe (Figure 4; see supplementary for significant parcels in the distance between Third Thumb and sensory feedback, and Third Thumb and D1).

**Figure 4:**
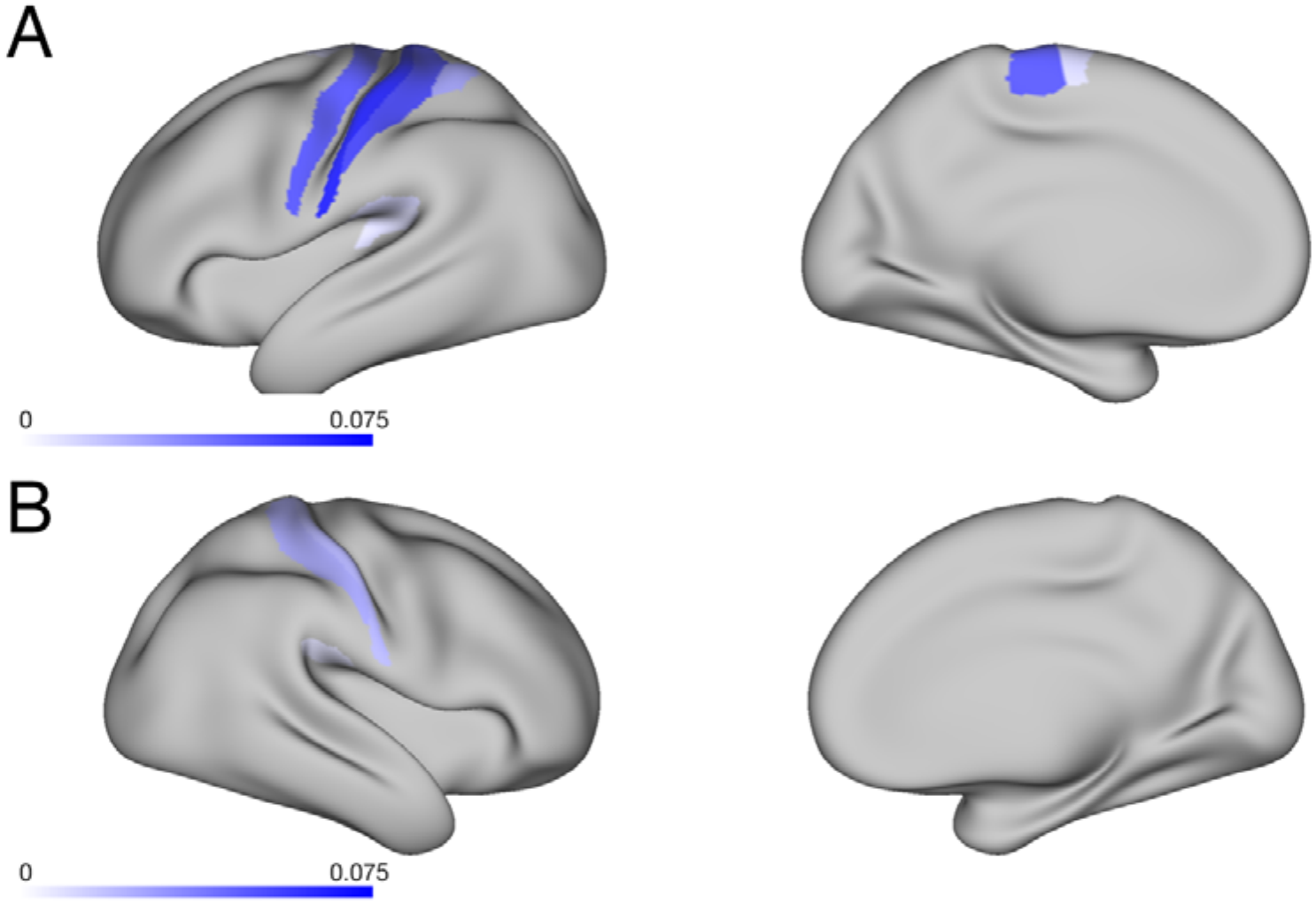
Experience-dependent differences in Third Thumb-Toe representational distance across the cortex. Cortical paracellations showing a significantly greater crossnobis distance between the Third Thumb and toe movement in the long-term compared with the short-term training group following permutation-based maximum-|t| FWER correction across parcels (p<.05). Surface colour indicates the mean difference in crossnobis distance between groups (long-term training – short-term training), with larger values (darker purple) reflecting greater Third Thumb-toe representational distance following long-term training. (A) Left hemisphere. (B) Right hemisphere.

There was a significantly larger distance between the Third Thumb and toe following long-term training across primary sensorimotor areas, extending the ROI analysis findings with significant parcels in left primary motor cortex (area 4) and bilateral primary somatosensory cortex (areas 1 and 2). Additional significant parcels were distributed across higher-order sensorimotor regions, including premotor and superior parietal cortex (left 6mp and 7PC), as well as higher-order somatosensory and parietal-opercular regions (left OP1, left RI and bilateral PFcm). This exploration indicates that the greater disassociation between the Third Thumb and the toe following long-term training extends beyond M1 to a broader cortical sensorimotor network.

## Discussion

There are long-held theories about the organisation of sensorimotor cortex, classically exemplified by the topographic organisation of the cortical homunculus (Penfield & Boldrey, 1937). Early neuroimaging work supported this view, characterising body representations through peaks of activity and regions showing preferential responses to individual body parts (Kleinschmidt et al., 1997; Rao et al., 1995). More recently however, looking beyond activity magnitude to the information carried across distributed patterns has revealed a more overlapping, ‘mosaic’-like organisation, with information about different body parts intermixed across cortical territories (Deo et al., 2026; Huber et al., 2020; Muret et al., 2022). Under this framework, representing an action does not necessarily require a selective, spatially discrete patch of cortex devoted to the effector performing it.

Our short-term experience results fit within this distributed view. Despite the absence of positive activity in the M1 hand area during Third Thumb control, its activity pattern was already reliably distinguishable from the biological thumb, toe movement and sensory feedback conditions. At the same time, PCM showed that this pattern was strongly dominated by the toe-related component. Thus, from initial experience, Third Thumb control already evokes a distinguishable neural state within M1, but one strongly anchored to the pre-existing, accessible biological signal used to control the device. This familiar motor signal may scaffold the rapid ability to use the Third Thumb from first exposure (Clode et al., 2024), without requiring dedicated cortical resources for the device itself.

Following prolonged experience however, there was a striking increase in M1 hand area activity during Third Thumb control. The lack of such activity in the short-term group establishes this recruitment is not required for Third Thumb use, yet the most prominent experience-dependent difference in our whole-brain analysis was centred on the sensorimotor hand knob. Given our extended work on the stability of body representation in primary sensorimotor cortex, even following profound changes in sensorimotor experience such as amputation (Kikkert et al., 2016; Schone et al., 2025; Wesselink et al., 2019), the emergence of such a marked and selective increase in hand area recruitment is striking. Such recruitment may be expected in higher-order regions reflecting increased planning, cognitive demands or contextual understanding associated with skilled Third Thumb use, but increased M1 activity is not a general signature of motor learning (Bassett et al., 2015; Berlot et al., 2020).

It is therefore tempting to interpret this increased recruitment as M1 ‘growing’ a representation of the extra digit - why else should activity in the hand area increase so selectively with experience? Indeed, neural changes to the body representation following use of supernumerary robotic fingers have previously been interpreted as evidence for functional reorganisation or incorporation of the artificial effector into the body representation (Hussain et al., 2017; Liu et al., 2022; Rossi et al., 2021; though please note that these studies did not disassociate the artificial effector from the biological sensorimotor signals supporting its use). Such an interpretation is also consistent with our previous finding that Third Thumb training reduced the representational distances between the biological fingers (Dowdall et al., 2026; Kieliba et al., 2021). Yet increased activity alone cannot establish that a dedicated Third Thumb representation has developed. It could equally reflect altered relevance of Third Thumb actions to hand function, increased engagement by their sensory consequences, or changes in how the constituent signals involved in device control are integrated. Multivariate analyses allow us to interrogate these possibilities more directly.

Following prolonged Third Thumb experience, its activity pattern becomes increasingly dissimilar from the controller toe movement, alongside a substantial down-weighting of toe-related information. Thus, although the toe remains the physical controller, the neural state associated with using it becomes increasingly differentiated from toe movement itself across a sensorimotor network.

Does this differentiation support the development of an independent Third Thumb representation? Partially, the increasing distance from the Toe suggests that Third Thumb control is becoming less strongly anchored to the biological resource through which it is being controlled. However, this differentiation does not provide sufficient evidence for an independent motor representation. The Third Thumb’s distance from the sensory feedback condition remains stable, while the relative weighting of this sensory component within the combined model increases substantially. Moreover, the Thumb’s observed representational structure can still be accounted for by a model combining the toe movement and sensory-feedback patterns, while adding D1 as an additional biological hand component does not improve this account. Our results therefore do not indicate the emergence of an independent representation, but a substantial re-weighting of the pre-existing sensorimotor resources that support Third Thumb control. This compositional organisation may extend a principle already evident in the biological motor system, where activity patterns for learned finger sequences can be largely explained by combinations of their constituent movement patterns (Yokoi et al., 2018). More broadly, this suggests that additional motor function can be accommodated through changes within an existing representational architecture, rather than requiring large-scale remapping or the allocation of dedicated cortical resources.

This sensory re-weighting may also help explain why M1 becomes so strongly recruited with prolonged use, where we also observed increased activity during passive Third Thumb movement. One possibility is that experience retunes M1 towards sensory signals that have become behaviourally relevant for Third Thumb control, fitting with evidence that somatosensory signals rapidly engage M1 and that their influence is shaped by behavioural context and task goals (Omrani et al., 2016; Pruszynski et al., 2014; Shelchkova et al., 2023). The intrinsic feedback generated on the hand is readily accessible and supports detailed perceptual judgements about Third Thumb interactions (Dowdall et al., 2026), and so may become increasingly utilised during skilled device use. Alternatively, the increased response may not reflect a specific role for this feedback in online control, but more general integration of Third Thumb-related sensory information into hand action representations through their repeated co-occurrence during device use (as seen for the Third Thumb in Dowdall et al., 2025; see also Dempsey-Jones et al., 2016; Ejaz et al., 2015). However, repeated co-occurrence alone cannot explain the increasing separation between the Third Thumb and Toe representations, which remain tightly coupled throughout training. Training may therefore also promote differentiation between two behaviours involving the same toe movement according to their functional role - moving the toe itself, or using that movement as the control signal for a distinct, hand-centred action, akin to the context-dependent differentiation observed during skill learning (Dash et al., 2025). The increasing contribution of sensory feedback may further help distinguish the consequences of these actions.

The relationship between the Third Thumb and biological hand actions provides complementary evidence that its representation may be shaped by the functional context in which the device is used. The Third Thumb activity pattern was consistently more similar to coordinated D1&Toe (its functional analogue) than D5&Toe (its topographic neighbour) movement, consistent with evidence that the representational geometry of the hand reflects everyday patterns of use (Ejaz et al., 2015), and that motor cortical organisation can reflect digit actions (Huber et al., 2020). We cannot tease apart if greater similarity is reflecting patterns of co-use (Dowdall et al., 2026; Kieliba et al., 2021), similarities in action kinematics, or a more abstract representation of general function. Nevertheless, these possibilities all point towards the statistics and structure of action as relevant for organising the Third Thumb within M1.

Finally, this continued grounding in existing sensorimotor channels raises an interesting question in light of behavioural generalisation. Third Thumb learning can transfer to a novel hand and controller mappings (Molina-Sanchez et al., 2026), suggesting that the learned skill is not irreversibly tied to the biological signals through which it was acquired. The increasing differentiation from the Toe is compatible with reduced dependence on this original controller, yet the increased contribution of hand-centred sensory information shows that the M1 representation itself is not body-independent. Prolonged training may therefore support a more flexible mapping between a higher-level Third Thumb skill and the sensorimotor resources used to implement it in a given context. M1 may reflect the current sensorimotor implementation of Third Thumb use, while transferable aspects of the learned skill depend on a more flexible action representation or control policy. Effector-independent representations have similarly been demonstrated for learned motor sequences (Wiestler et al., 2014), although whether a comparable mechanism supports generalisation of complex collaborative devices such as the Third Thumb remains to be established. Determining where and how such mapping flexibility is represented will require experiments that directly manipulate the hand and controller used during imaging.

In conclusion, prolonged experience with an additional robotic effector changes both the recruitment and organisation of existing M1 resources. Third Thumb control is distinguishable from early experience but initially remains strongly anchored to that familiar motor signal of its biological controller. With prolonged training, this relationship weakens while sensory consequences arising on the hand become increasingly prominent. Rather than requiring a dedicated cortical territory for the artificial effector, these findings suggest that M1 can support an expanded motor repertoire by re-weighting the pre-existing sensorimotor resources through which the new action is controlled and sensed.

## Materials and Methods

### Participants

In total, 26 short-term training and 30 long-term training participants took part in the study. Two long-term participants were excluded because their scanning paradigm did not include the D1-only movement condition. Prior to data inspection, a further four short-term and six long-term participants were excluded due to acquisition-related issues: technical problems (*n*=5), participant illness or discomfort (*n*=4), or insufficient scanning time (*n*=1). The relatively high incidence of technical issues reflects the complexity of the experimental setup. Although participant illness or discomfort was not reported as being directly related to the paradigm, the MRI task was performed at the end of a ∼1.5-hour scanning session and, for the long-term group, following at least seven days of training, which may have contributed to reduced task tolerance.

The final dataset for analysis comprised of 22 short-term training participants (Age *M*=25.45 (*SD*=5.67), 15 female) and 22 long-term training participants (Age *M*=25.14 (*SD*=4.04), 13 female). There were no significant differences in the age (*t*(42)=.214,*p*=.831,BF_10_=.303) and gender (χ*^2^*(1,N=44)=.393,*p*=.531,BF_10_=.418) of the groups. All participants were right-handed. The study was approved by the Cambridge Psychology Research Ethics Committee (PREC: 2022.068) and the UCL Research Ethics Committee (project ID number 12921/001). Participants were recruited via online advertisements on the University College London, MRC Cognition and Brain Sciences Unit and University of Cambridge SONA participant pools. Participants had no known neurological and/or sensorimotor disorders, had no previous experience with the Third Thumb, and had no more than five years of musical experience, and had not acquired any musical training within the last five years.

A between-groups design was used to match task-specific experience with the Third Thumb in the scanner. By avoiding repeated exposure to the fMRI task, both groups encountered the scanner task specific movements with comparable novelty. This ensured differences between the short- and long-term experience groups primarily reflected their prior experience with the Third Thumb, rather than reflecting differences in familiarity or practice with the task itself.

### The Third Thumb

The Third Thumb (Dani Clode Design, Cambridge, UK) is a robotic extra digit that attaches to the ulnar side of the right hand (Figure 1A) and is operated through pressure sensors placed underneath the big toes. The Third Thumb has a possible two degrees of freedom (DoF) that allow a corresponding proportional control; applying pressure to the right big toe causes a flexion/extension movement; whilst pressure on the left causes an adduction/abduction movement (see further details in Clode et al., 2024 and Kieliba et al., 2021).

An adapted Third Thumb was used in the MRI scanner. A non-ferromagnetic Third Thumb set-up (Figure 5) was built in order to safely use the device in the MRI scanning environment. The control sensor was fixed to an adjustable footrest slotted into the base of the scanner bed and safely connected to the microcontroller in the control room. While two DoF are possible with the MRI Third Thumb, only the flexion/extension DoF was enabled for the task, which minimised head movements and ensured task ease given the lack of vision.

**Figure 5:**
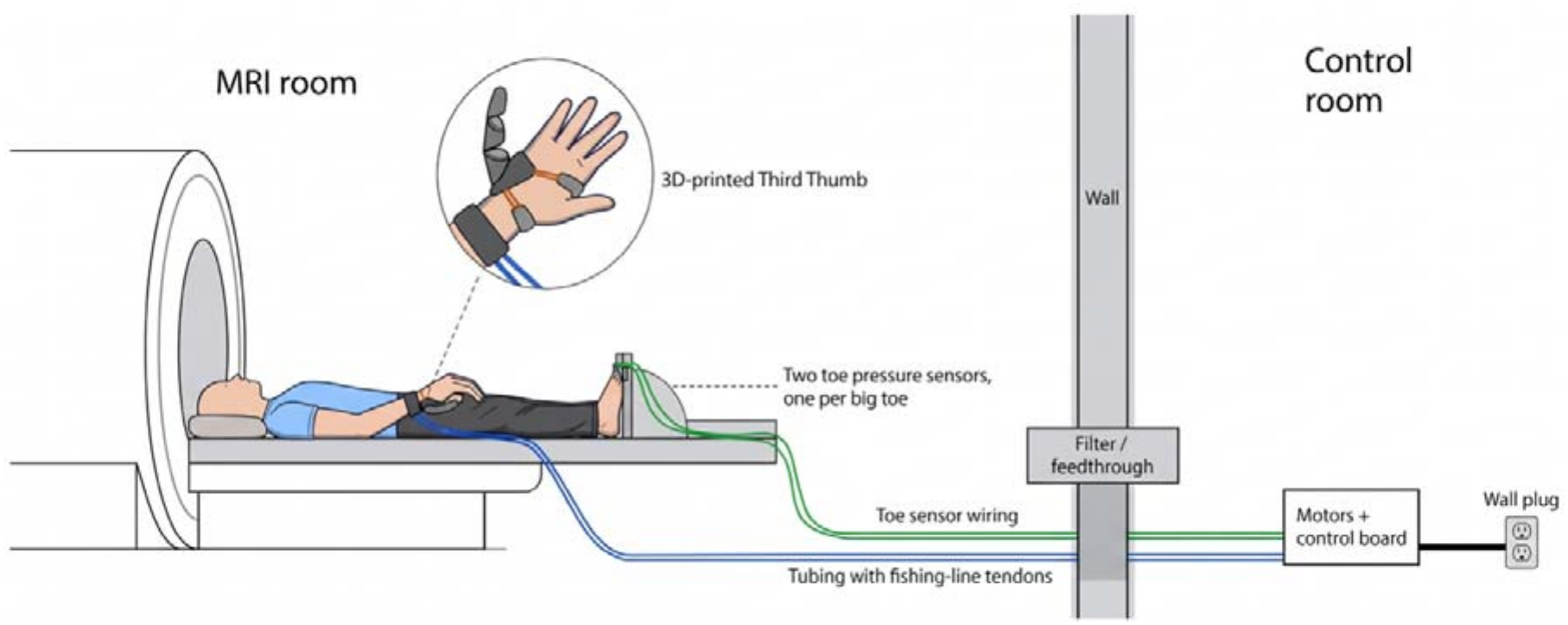
Third Thumb MRI setup. In the scanner, the Third Thumb hand piece was fitted to the participants’ right hand. It was controlled by applying pressure by their right big toe to a pressure sensor positioned on a footrest fixed underneath the participants’ feet. This controlled the one degree of freedom used in the MRI task. An object attached to the hand provided a fixed tapping target, and the hand was positioned comfortably across the body while participants lay in the scanner. A serial cable then passed through the radiofrequency filter panel, connecting the toe pressure sensor to the Arduino microcontroller within the control room. The Third Thumb movement was then controlled via servomotors within the control room, connected to the Thumb through non-ferromagnetic teflon tubes.

Given the one DoF MRI task and only limited training time beforehand, short-term experience participants trained with a one DoF Third Thumb. Long-term experience participants training with the two DoF Third Thumb.

### Training Tasks

Short-term training participants were first familiarised with the Third Thumb for 90 minutes. Within the session they were trained on a series of tasks designed to develop basic motor control of the device. These tasks included: Coordination – between the biological fingers and Third Thumb: opposing the tips of the fingers to the tip of the Thumb; Individuation – of the Third Thumb (picking up, transporting and repositioning tape rolls) while keeping the biological hand occupied; Collaboration – between the biological hand and Thumb: lifting and transporting wooden bars to build a tower. Each of these tasks was administered for approximately 10 minutes (for further details see Supplementary Materials).

Long-term training participants underwent seven days of training, designed to allow acquisition of different motor skills with the Thumb – including the individuation, collaboration and coordination experienced by the short-term training group. The first and last day of training was supervised in-person, whilst the middle five consecutive days of training were completely remotely and semi-supervised (approximately two hours of training per day). For details on the training for the long-term training participants, see Molina-Sanchez et al., 2025. Before training, participants underwent two sessions with the Third Thumb (a familiarisation, then pre-training session) and underwent a post-training session after the seven days.

### Functional Magnetic Resonance Imaging (fMRI)

#### Acquisition

MRI acquisition occurred at the Functional Imaging Laboratory (FIL) (London, UK) for the short-term training group, and at the MRC Cognition and Brain Sciences Unit (CBU) (Cambridge, UK) for the long-term training group. Identical acquisition parameters were used at both sites.

MRI images were acquired using a 3T Prisma Fit MRI scanner (Siemens, Munich, Germany) using a 32-channel head coil. For functional image acquisition, a multiband T2* -weighted pulse sequence with a between-slice acceleration factor of 4 and no in-slice acceleration was used. A voxel size of 2mm isotropic was used, and a repetition time (TR) of 1500 ms. Acquisition sequence parameters consisted of an echo time (TE) of 35 ms, flip angle of 70° with 72 transversal slices per volume. For the T1-weighted image acquisition sequence, to acquire the structural image we used a voxel size of 1mm isotropic and TR of 2250 ms, with sequence parameters consisting of a 3ms TE, flip angle of 9°.

#### Functional Task

Participants performed a short task familiarisation before going into the scanner, performing the task for approximately 10 minutes lying down on a mock MRI bed. Once on the scanner bed, the Third Thumb was fitted to the participants’ hand, followed by a custom 3D printed ergonomic object that was strapped around the hand. With the support of foam pads, the hand was placed at approximately 90° across the participants’ body, ensuring the hand and Third Thumb had full range of motion to tap on the object. The footrest was adjusted so the participant could press easily on the pressure sensor that controlled the Third Thumb with their right big toe, producing Third Thumb movement and minimal head movements.

During the scan, participants performed a motor tapping task. All hand movements involved tapping on a hand-held object. Conditions were presented visually. Participants performed six separate conditions: (i) Tap the Third Thumb (ii) Move their right toe (iii) Passively experience the Third Thumb moving autonomously to tap on the object (iv) Tap D1 (v) Tap D1 while also moving their right toe (vi) Tap D5 while also moving their right toe. See full condition details above in Results.

In conditions involving the toe with no Third Thumb, the pressure sensor was temporarily turned off but participants were instructed to apply the same pressure with their toe in all relevant conditions.

Due to constraints in the time available for the task within the scanning session, the groups differed in their specific block and run parameters. Participants completed three (long-term training) or four (short-term training) runs of the task, with 170 (long-term) or 207 (short-term) volumes.

Conditions were presented visually (via Psychopy, minimum version 2021.2.1). The condition name appeared in white for two seconds, then flashed green three (long-term training) or four (short-term training) times and in red once for the last movement – resulting in four (long-term training) or five (short-term training) movement cycles per block; with six or eight seconds of movement. Movements were performed at a frequency of approximately 0.5 Hz. To avoid visual contamination, the length of the characters on the screen were matched by adding ashes (#) when necessary, and conditions were projected onto a pink noise black background. Each block was repeated four times per run, with an additional six rest blocks. Conditions were pseudo-randomly distributed, such that each condition was equally preceded by all other conditions. To confirm correct movements were made at the instructed times, task performance was visually monitored online through the video output from the scanner cameras.

#### Pre-processing and first-level analysis

All MRI data pre-processing and analyses were carried out using FMRIB Software Library (Jenkinson et al., 2012; FSL, version 6.0.5) as well as in-house scripts written in MATLAB (version R2020b), Python (3.13.0), and R (4.2.2). For multivariate analyses, we used the MATLAB RSA Toolbox (Nili et al., 2014) in addition to within-lab toolbox extensions (Wesselink & Maimon-Mor, 2017) and the Python PCM Toolbox (Diedrichsen et al., 2011, 2018), in addition to custom Python scripts.

To ensure that the functional scans were well aligned, a midspace was calculated between the runs for each participant. This represents the average space where the images are minimally reoriented. Each scan was aligned to this midspace using FLIRT (Jenkinson et al., 2002; Jenkinson & Smith, 2001), with structural image alignment optimised using Boundary-Based Registration.

Functional data was primarily pre-processed using FSL-FEAT. Motion correction was performed using FMRIB’s Linear Image Registration Tool (MCFLIRT (Jenkinson et al., 2002)), followed by brain extraction using the Brain Extraction Tool (BET (Smith, 2002)). The data then underwent temporal high-pass filtering, with a cutoff of 90 seconds, and lastly spatial smoothing with a 3 mm full width at half maximum (FWHM) Gaussian kernel. Field maps were also used for field unwarping.

The resulting functional images were used for all subsequent first-level GLM analyses. A voxel-based general linear model (GLM) was implemented in FSL-FEAT to identify activity patterns related to each movement condition. For each run, six regressors were created, one for each condition (Third Thumb, Toe, Feedback, D1, D1&Toe, D5&Toe). Regressors were convoluted with a double-gamma function to account for the delayed BOLD response. Additional first derivative regressors were included to capture temporal variations in the BOLD signal, along with six motion parameters to estimate head movements. Additionally, we used motion scrubbing, adding regressors of no interest to exclude any volumes with excessive motion (framewise displacement larger than 0.9 (Power et al., 2013)). In the model, 6 contrasts were set up, namely each condition vs rest.

For each participant, the three (long-term) or four (short-term) contrast images for each movement condition obtained from the first-level analysis were combined across runs using a fixed-effects second-level analysis in FSL FEAT. The resulting contrast images were then normalised to MNI152 standard space.

Group analysis was then performed using the second-level contrast images. The Third Thumb activity was contrasted between the long and short-term experience groups to determine experience-dependent differential activity (Figure 2C). This used a random-effect model in FSL (FLAME 1) and a cluster correction to control for multiple comparisons using a Z-threshold of 2.6 (corresponds to a two-tailed p value of approximately 0.01) and a cluster p-threshold of .05. The resulting thresholded map was projected onto a standardised surface for visualisation purposes only with 5mm smoothing applied.

#### Region of Interest (ROI) Definition

Analyses were focused on the M1 hand area, with supplementary analyses on the S1 hand area. First the anatomical ROIs were defined in the left hemisphere based on probabilistic cytoarchitectonic maps (Wiestler & Diedrichsen, 2013) and aligned to the average Freesurfer surface (Fischl, 2012; Fischl et al., 2008). The hand region of primary motor cortex (M1) was defined as Brodmann area (BA) 4, 2.5 cm above and below the hand knob (Yousry et al., 1997). S1 was similarly defined as the hand-related aspect of BA 1,2, and 3. However, we note that, given the probabilistic nature of these masks, the dissociation between S1 and M1 is only an estimate. Structural T1-weighted images were used to reconstruct the pial and white-grey matter surfaces using Freesurfer. The anatomically defined ROIs were projected onto the individual brains via the reconstructed individual anatomical surfaces.

To identify the voxels most selective to hand representations, while ensuring the same ROI size across participants, the anatomical ROIs were spatially restricted using a contrast of coordinated movement of the index finger (D2), Third Thumb and toe (vs baseline) from a different task. Within each anatomical ROI and participant, the top 300 most active voxels were selected. For the M1 foot area, the top 150 most active voxels were selected.

#### Representational Similarity Analysis

To explore the informational content in each condition, we used multivariate representational similarity analysis (RSA; Kriegeskorte et al., 2008)). For each participant and each run, the beta weights estimates were extracted from the first-level analyses for each condition vs rest within our hand area ROI. These were prewhitened using the GLM residuals, and the cross-validated Mahalanobis (crossnobis) distance (Nili et al., 2014) was calculated between the patterns for each of the conditions. The distances were calculated using each possible imaging run pair combination within a session and then averaged the distances across all run pairs. This distance provides a measure of dissimilarity in the activity patterns elicited in each movement condition. The expected distance is zero if two patterns are not significantly different from each other.

#### Pattern Component Modelling

To determine whether the Third Thumb representation could be explained by a recombination of its constituent motor and sensory components, we used pattern component modelling (PCM; Diedrichsen et al., 2018), implemented in PcmPy. PCM characterises the representational structure of multivariate activity patterns through their second-moment matrix, (G). For (K) experimental conditions measured across (P) voxels: 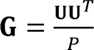 where (U) is the condition-by-voxel matrix of activity patterns. The diagonal elements of (G) describe the pattern variance associated with each condition, whereas the off-diagonal elements describe the covariance between condition-specific patterns. Thus, G specifies the variance of each condition-specific activity pattern and the covariance between patterns. These quantities determine their relative similarities and pairwise distances. The model therefore predicts the relationships among condition-specific multivoxel patterns, rather than the expected response of each individual voxel.

For each participant, run-specific voxel-wise activity estimates from the M1 hand area ROI were entered for the toe movement, sensory feedback and Third Thumb conditions, with imaging run specified as the partition variable. The three models constructed made different predictions about the (3X3) second-moment matrix relating these conditions, with diagonal entries describing the variance of each condition-specific pattern and off-diagonal entries describing the covariance between patterns (Figure 6).

**Figure 6:**
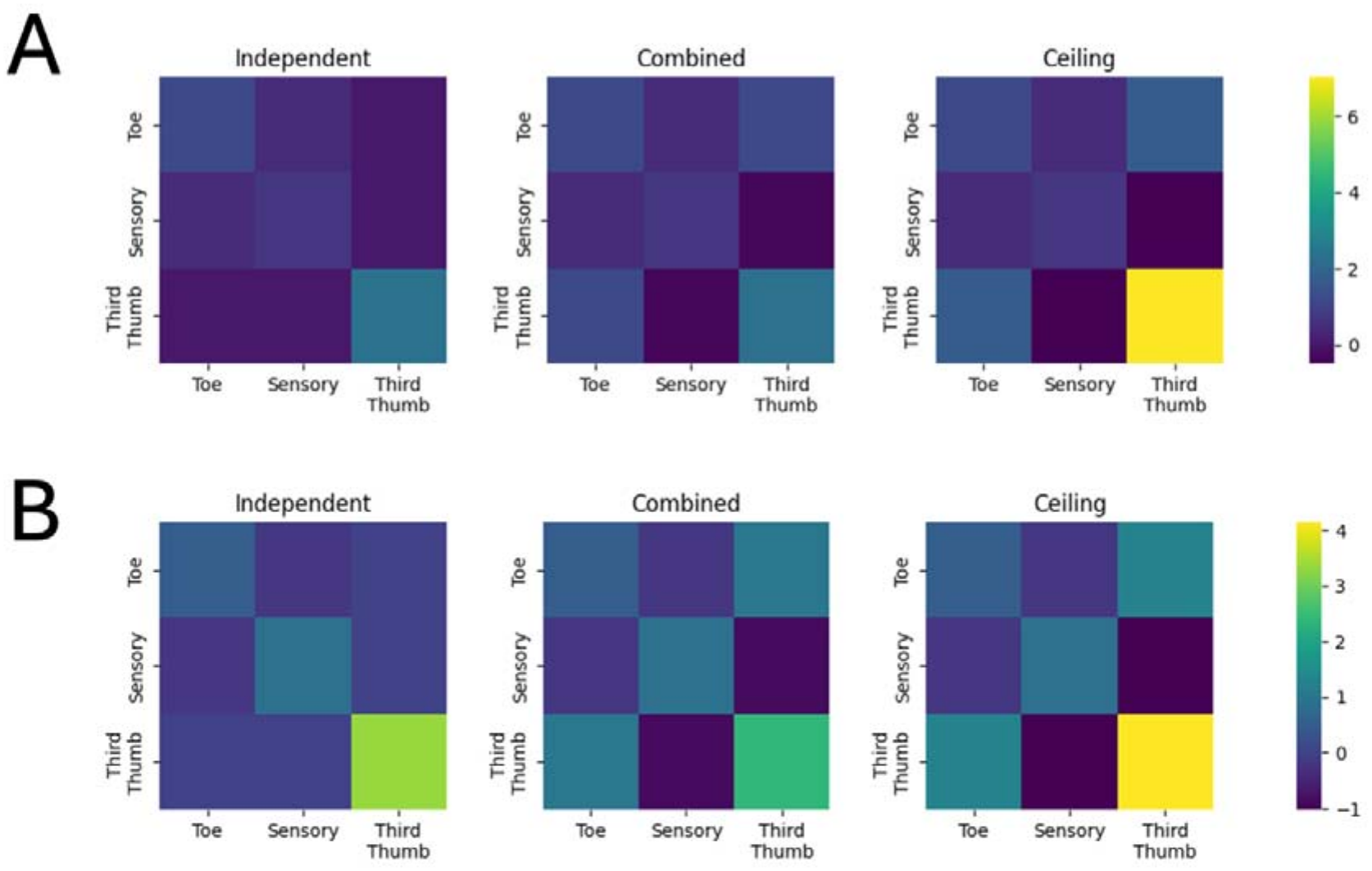
Model-predicted second-moment matrices for the PCM models. Predicted G matrices after fitting the model parameters separately for the short-(top) and long-term (bottom) experience groups. Rows and columns correspond to the toe movement, sensory feedback and Third Thumb conditions. Diagonal elements represent the pattern variance associated with each condition, while off-diagonal elements represent the covariance between condition-specific activity patterns. The independent model constrains the Third Thumb pattern to be orthogonal to the toe and sensory feedback patterns; the combined model constrains the Third Thumb pattern to a weighted linear combination of these two constituent patterns; and the noise-ceiling model freely estimates the variance–covariance structure among all three conditions. Colour indicates the fitted variance (diagonal) or covariance (off-diagonal) value, with separate colour scales for the short- and long-term groups.

The combination model tested whether the Third Thumb pattern lay within the representational space spanned by the toe movement and sensory feedback patterns. The variance and covariance of these two component patterns were freely estimated, while the Third Thumb pattern was constrained as:

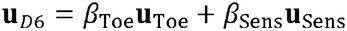

This model contained five representational parameters: two parameters describing the variances of the toe movement and sensory feedback patterns, one parameter describing their covariance, and two parameters describing their contributions to the Third Thumb pattern. Because no additional Third Thumb-specific component was included, the model predicted that all reliable structure in the Third Thumb representation could be expressed as a linear recombination of the two constituent patterns.

The independent model instead assigned the Third Thumb an independent pattern component, constraining its covariance with both constituent patterns to zero, meaning the pattern was orthogonal to both the toe movement and sensory feedback patterns.

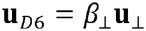

A fully flexible free model then freely estimated all variances and covariances among the three conditions, subject only to the requirement that G represented a mathematically valid second-moment structure. This model was taken as the lower noise ceiling estimate.

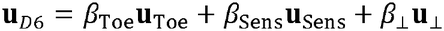

A true null model was also included, with the second-moment matrix fixed to zero (G=0), representing the absence of reliable condition-related signal across all three conditions. Unlike the independent model, this model allowed no representational variance or covariance, providing a baseline against which model performance could be evaluated.

Models were fitted separately within the short- and long-term experience groups using leave-one-participant-out cross-validation. In each fold, the representational parameters were estimated jointly from all remaining participants and evaluated in the held-out participant. Participant-specific signal-scaling and noise parameters were estimated separately, allowing overall response magnitude and noise levels to vary across participants. Residual noise was modelled using a block-plus-independent covariance structure, allowing activity estimates obtained within the same imaging run to share noise while also estimating independent residual variance for each observation.

Model performance was quantified using the cross-validated log-likelihood of the held-out participant’s voxel-wise activity data, with higher values indicating a better account of the participant’s representational structure. When visualising model performance, log-likelihoods were expressed relative to the true null model by subtracting its log-likelihood for each participant. For direct model comparisons, differences in log-likelihood were calculated. The combination and independent models were compared to test whether the Third Thumb was better explained as a recombination of its constituent patterns than as an unrelated representation. The combination model was then compared to the noise ceiling model to assess model adequacy. A combination model that performed below the free model would indicate that reliable representational structure remained unexplained. Participant-level log-likelihood differences were tested against zero.

The fitted toe movement weighting and sensory feedback weighting parameters were extracted from the combination model to characterise how the two constituent patterns contributed to the Third Thumb representation. These coefficients were unconstrained: they could take positive or negative values and were not required to sum to one. To assess their relative weighting, each coefficient was normalised by the sum of the toe movement and sensory feedback coefficients. This yielded proportional weightings that summed to one, allowing the relative weighting of sensory feedback and toe movement to be compared between groups.

As a biological comparison, the combined model analysis was replicated using toe movement, biological thumb movement and their coordinated D1&Toe condition. The combination model predicted:

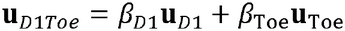

The corresponding independent and free models were fitted using the same cross-validation procedure, providing a benchmark for whether the representation of a familiar coordinated biological action could be described as a recombination of its constituent movement patterns.

The final model tested whether biological thumb movement explained additional structure in the Third Thumb representation beyond toe movement and sensory feedback. The extended model included toe movement, sensory feedback and D1 as explanatory patterns and constrained the Third Thumb representation as:

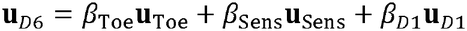

The covariance structure among the three explanatory patterns was freely estimated. To isolate the additional contribution of D1, a reduced model was constructed using the same four conditions and covariance structure, but with the D1 weighting fixed to zero (β D1=0).

### Whole Cortex Parcellation Analysis

As an exploratory analysis, to examine experience-dependent differences in representational geometry beyond the predefined ROIs, we extended the RSA across the cortex utilising the parcellations defined by the Human Connectome Project multimodal parcellation (HCP-MMP1.0; Glasser et al., 2016). The 360 cortical parcels were mapped from the standard fs_LR surface to each participant’s native cortical surface and subsequently converted to the participant’s anatomical T1-weighted image using ribbon-constrained surface-to-volume mapping. Within each parcel, crossnobis distances were calculated using the same prewhitening and cross-validation procedure described above, focusing on the distances between the Third Thumb and toe movement, sensory feedback, and D1.

For each distance, the long- and short-term training groups were compared within each parcel using student’s independent-samples t-test. Multiple comparisons across parcels were controlled for each contrast of interest using a permutation-based maximum-statistic procedure. Group labels were randomly permuted at the participant level 100,000 times while preserving group sizes, and the maximum absolute t-statistic across parcels was retained for each permutation. Observed parcel-wise statistics were compared against this null distribution to obtain two-sided family-wise error rate (FWER)-corrected p-values, calculated separately for each representational distance.

### Controlling for Between-Group Differences

The short- and long-term training group were acquired at different imaging sites, meaning site was confounded with group. We therefore took several steps to minimise and assess the potential contribution of differences in acquisition, task protocol and data quality to between-group effects. Both datasets were acquired using 3T Siemens Prisma Fit MRI scanners with 32-channel head coils and identical structural and functional acquisition parameters. Participants also performed the same six experimental conditions using the same one-degree-of-freedom MRI-compatible Third Thumb setup and movement frequency.

The same preprocessing and first-level analysis pipeline was applied to both datasets. Given the motor demands of the task, additional assessments were performed to ensure head motion did not differ systematically between groups. Six motion parameters were included as nuisance regressors in each first-level GLM, and volumes exceeding a framewise-displacement threshold of 0.9 were additionally modelled using motion-scrubbing regressors. The proportion of volumes per run identified as motion outliers did not differ significantly between groups (BF_10_ .268). The mean proportion of outlier volumes was 0.3% for the short-term group and 0.7% for the long-term group. Motion outliers were detected in 22.6% and 11.1% of runs, respectively.

As an additional sensitivity analysis, the long-term training dataset was denoised using MELODIC and FSL FIX with a study-specific classifier to assess whether movement-related artefacts could account for the observed effects. Applying this additional denoising did not alter the principal univariate, RSA or PCM results (see *Supplementary Materials*). All analyses reported in the main manuscript therefore use data performed without FIX.

Reliability of the multivoxel activity patterns was also assessed across imaging runs as a measure of noise ceiling. For each participant and experimental condition, Pearson correlations were calculated across voxels between the condition-specific activity patterns from every pair of independent runs. Correlations were averaged within participant to provide a measure of run-to-run pattern reliability. Pattern reliability was comparable across groups (no difference for tested patterns of D1 (BF_10_=.318), D1 and Toe (BF_10_=.317) nor Third Thumb (BF_10_=.319)), providing no evidence for a systematic difference in the stability of the multivoxel patterns between datasets.

Finally, the principal analyses were designed to emphasise condition-specific and within-participant representational effects rather than absolute signal differences between datasets. Univariate group effects were assessed primarily through Group X Condition interactions, using stable within-participant biological conditions as reference points. RSA quantified representational distances from prewhitened activity estimates using cross-validation across independent imaging runs, while PCM modelled the representational structure separately within each group and included participant-specific signal-scaling and noise parameters. Together, these approaches reduce sensitivity to global differences in signal magnitude or measurement quality between sites.

### Statistical Analysis

All statistical comparisons were performed in JASP (version 0.19.1). Throughout, parametric statistics were used when there were no violations of test assumptions (normality assessed by Shapiro-Wilk values and visual inspection, homoscedasticity assessed by Levene test values). If there were violations, when possible, the equivalent non-parametric test was used. All null results of interest were followed up using Bayesian statistics (Cauchy prior width r=0.707) to obtain a Bayesian factor (BF_10_) to allow interpretation based on the strength of support for the null hypothesis. We used a threshold of BF<1/3 as sufficient evidence in support of the null hypothesis, BF>3 as sufficient evidence in support of the alternative hypothesis, and 1/3<BF<3 as inconclusive evidence (Wetzels et al., 2011). Where Bayesian factors could not be obtained directly for effects tested using linear mixed-effects models, an equivalent t-test or ANOVA was performed to obtain an approximate BF_10_ for the corresponding effect (indicated by ≈).

Wherever possible, we employed linear mixed models (LMM), with a random intercept for participant. We used the Satterthwaite testing method and Satterthwaite degrees of freedom approximation for all follow-up tests. As fixed effects, models accounted for group and condition, in addition to model for the PCM analysis. If the model could not converge, a repeated measures ANOVA was used instead.

For any simple comparisons where a LMM would not be appropriate, we used one-sample t-tests to assess if values were significantly different from zero (including average z-stats and RSA distances), paired samples t-test for within-subjects comparisons (including when comparing the PCM model fits and weightings), and independent samples t-tests for between-subjects comparisons (including when comparing average z-stats, RSA distances and PCM weightings). A Wilcoxon signed-rank test (within-subjects) or a Mann-Whitney test (between) was used if assumptions were violated.

## Supplementary Materials

**Figure S1:**
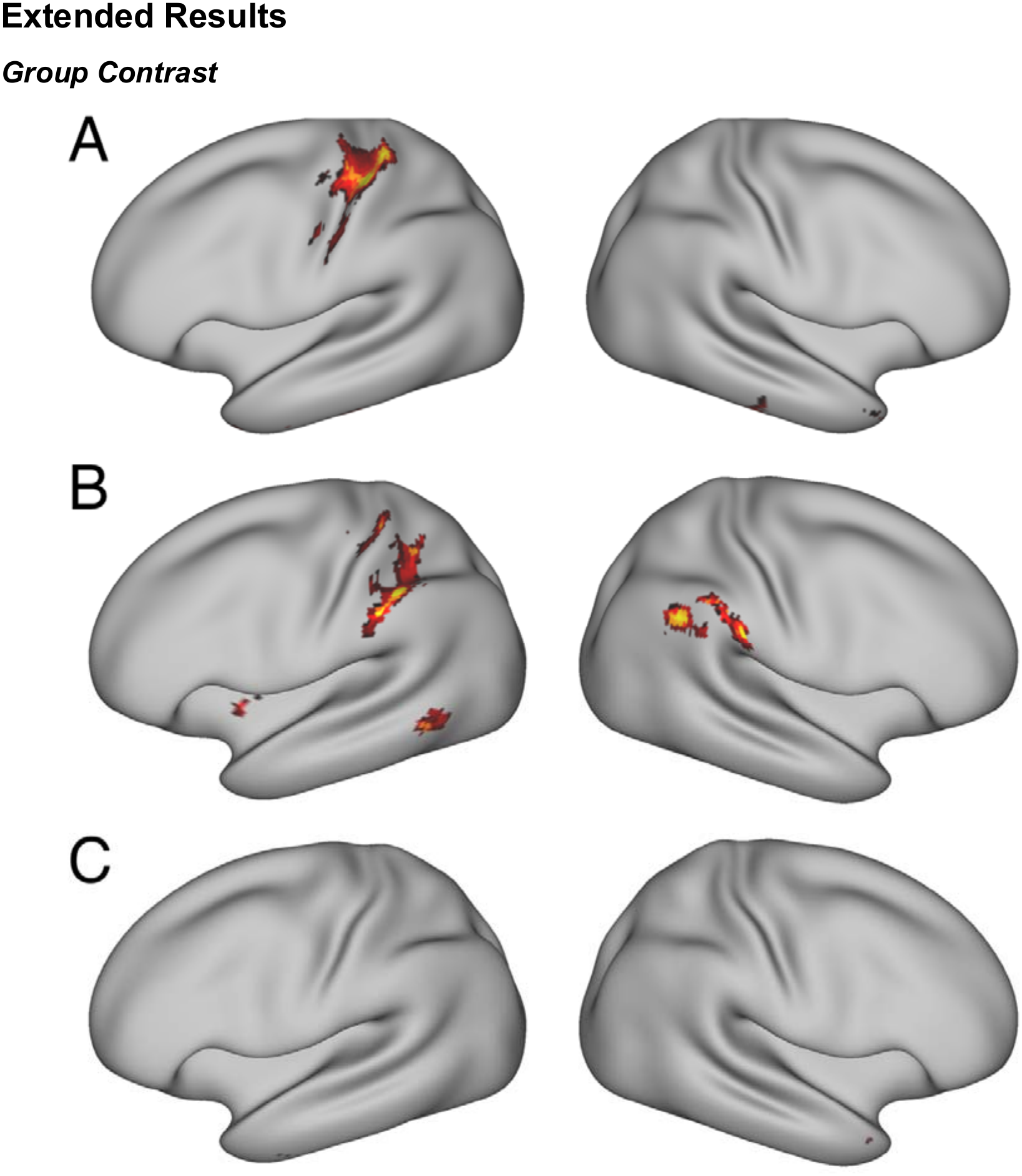
Whole-brain group contrasts. Whole-brain contrast of activity between the long- and short-term experience groups during the (A) Third Thumb movement condition (B) sensory feedback (passive Third Thumb movement) condition and (C) toe movement only condition. (A) Greater activity following long-term experience centred around the sensorimotor hand knob, with an additional small cluster in inferior temporal gyrus (B) Greater activity following long-term experience in S1 (centred around the sensorimotor hand knob) and in anterior interparietal sulcus, as well as the lateral occipital cortex (left hemisphere). Additional clusters of greater activity in inferior parietal lobe (right hemisphere) (C) One small cluster of greater activity following long-term experience in inferior temporal lobe. Positive values indicate greater activity in the long-term relative to the short-term group. The map is displayed at p<.01, cluster-corrected at p<.05.

**Figure S2:**
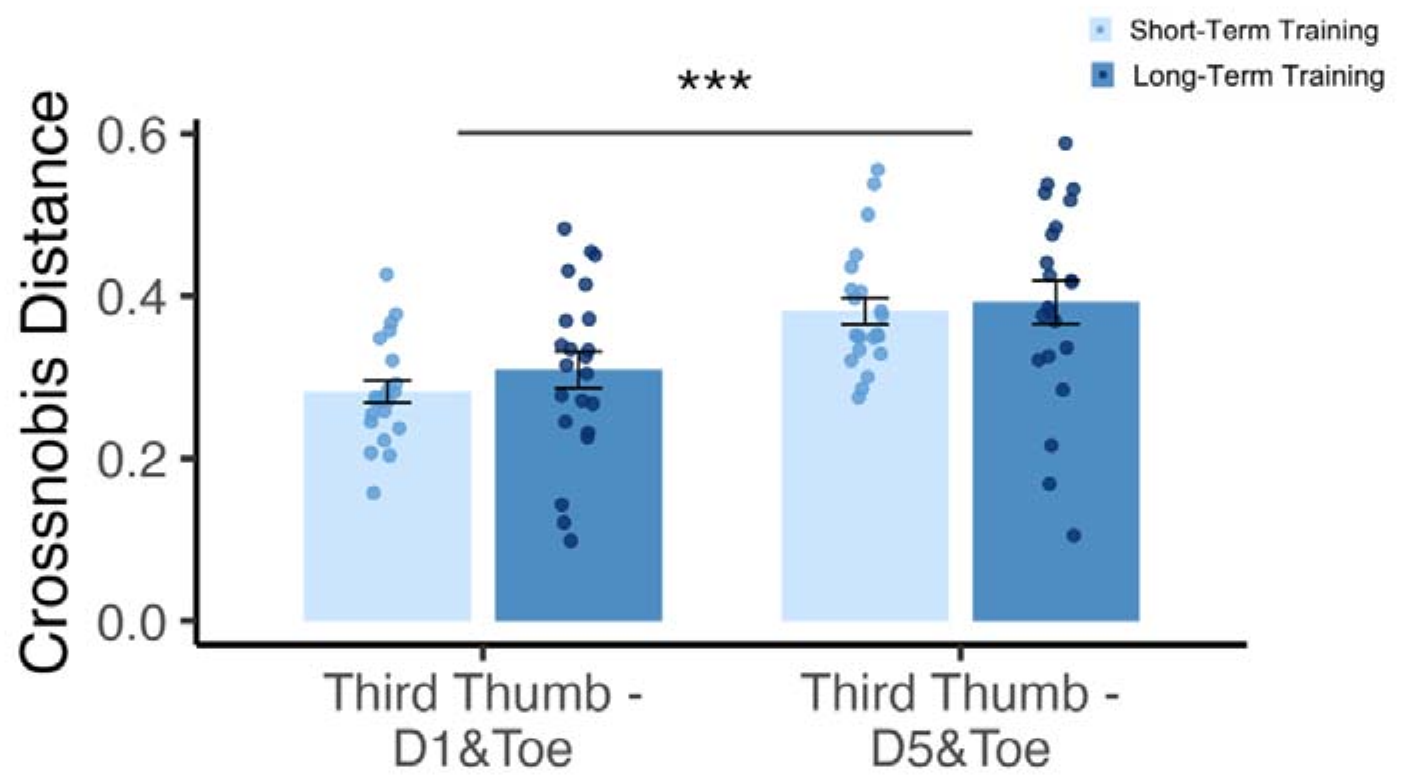
Representational similarity analysis within the M1 hand area. Crossnobis distances quantify the dissimilarity between the Third Thumb activity pattern and the biological thumb and toe movement (D1&Toe), and the little finger and toe movement (D5&Toe). The Third Thumb-D1&Toe distance was significantly smaller for both groups.

**Figure S3:**
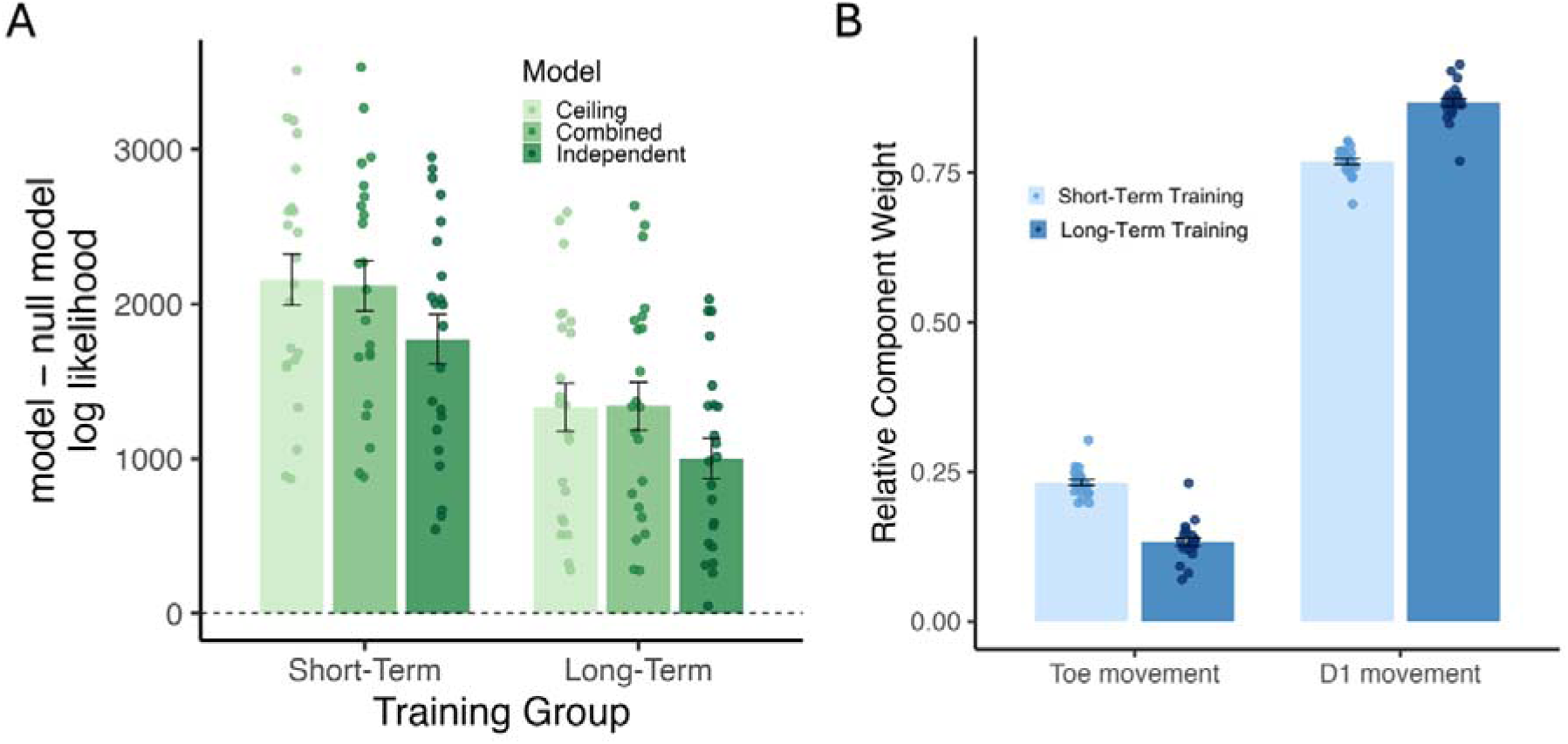
Pattern component modelling tested whether the D1&Toe pattern could be explained as a weighted combination of its constituent toe movement and D1 movement. (A) Cross-validated log-likelihoods for each model (independent, combined and noise ceiling) are subtracted from a null model (for visualisation only), with higher values indicating better model fit. (B) Relative component weightings estimated from the combined model.

### Primary Somatosensory Cortex (S1) Hand Area

#### Univariate Activity

Given the whole-brain contrast broadly identified the sensorimotor hand knob as the area of increased activity when controlling the Third Thumb in the long-vs short-term group, we replicated the core analyses from the main manuscript in the S1 hand area. Please note that, given the probabilistic nature of these masks, the dissociation between S1 and M1 is only an estimate.

One participant had negative activity across all biological finger conditions (D1, D1&Toe and D5&Toe) in the S1 hand area. They were therefore removed from all S1 hand analyses.

Contrary to M1, all of our core conditions significantly activated the S1 hand area: (Short-term group: D1 (*t*(21)=11.18,*p*<.001), Toe (*t*(21)=4.328,*p*<.001), Sensory feedback (*t*(21)=3.949,*p*<.001), Third Thumb (*t*(21)=4.377,*p*<.001); Long-term group: D1 (*t*(20)=11.52,*p*<.001), Toe (*t*(20)=3.583,*p*<.001), Sensory feedback (*t*(20)=5.874,*p*<.001), Third Thumb (*t*(20)=8.665,*p*<.001); Figure S2). While there was not a significant difference in D1 activity between groups (*t*(41)=1.816,*p*=.077,BF_10_=1.102), the inconclusive bayes factor meant all further group comparisons explored interaction terms relative to D1.

Within the S1 hand area, there was no difference in toe movement activity between the training groups (*F*(1,41)=.414,*p*=.523). There was a significant interaction for the sensory feedback (Group X Condition: *F*(1,41)=5.601,*p*=.023), however the follow-up contrast did not reach significance (short-vs long-term: *t*(79.61)=1.385,*p*=.170). We also found a significant interaction for the Third Thumb activity between-groups (*F*(1,41)=5.592,*p*=.023), but again the contrast did not reach statistical significance (*t*(79.83)=1.781,*p*=.079). These results echo the univariate results from M1, but with lower statistical clarity.

**Figure S4:**
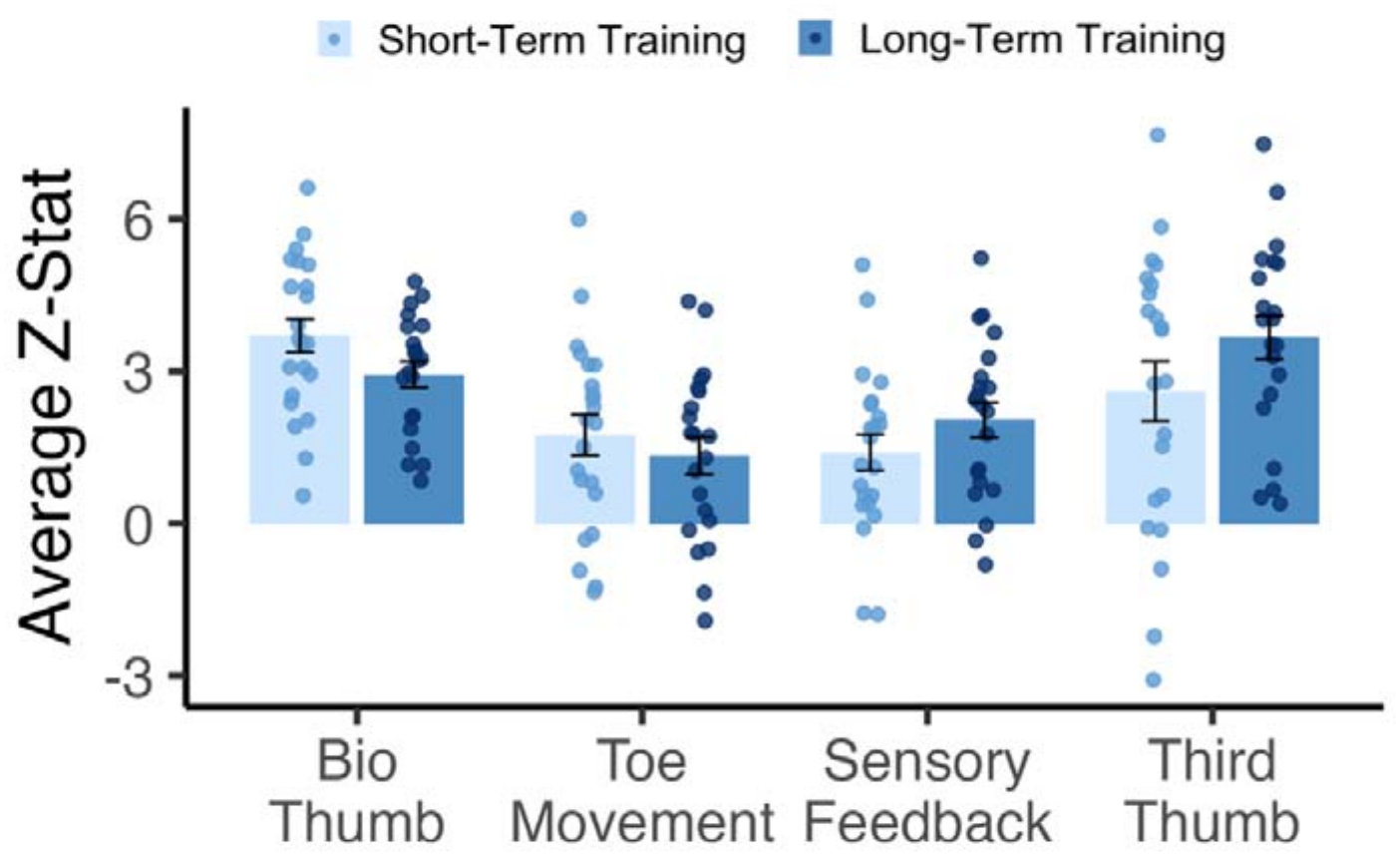
Univariate Activity in the S1 hand area. Average activity within the S1 hand-area ROI during biological thumb (D1), toe movement, passive sensory feedback and active Third Thumb movement following short- and long-term Third Thumb experience. All four conditions significantly activated the S1 hand area in both groups (p < .001). Toe movement activity did not differ between groups. Bars show group means and individual dots represent participants.

#### Representational Similarity Analysis

For the short- and long-term groups, all distances between the Third Thumb and each condition were significantly above zero (*p*<.001 throughout).

When considering if Third Thumb movement was represented more similarly to biological function (coordinated D1 and Toe movement) or anatomical position (coordinated D5 and Toe movement), we found it was consistently more similar to D1&Toe across groups (*F*(1,42)=50.009,*p*<.001). Between groups, the distance between D1 and the Third Thumb was not significantly different (*t*(41)=.184,*p*=.855,BF_10_=.304). This was replicated when taking the distance between the passive sensory feedback and D1&Toe vs D5&Toe: *F*(1,41)=33.995,*p*<.001, where the feedback was again more similar to D1&Toe across groups (not significantly different between; *t*(41)=.417,*p*=.679).

There was no significant difference in the distance between the Third Thumb and D1 (*t*(41)=.931,*p*=.357,BF_10_=.425), but with the inconclusive bayes factor, all further distances were still assessed in relation to the Third Thumb to D1 distance. The distance between the Third Thumb and toe movement was significantly greater following long-term experience compared to short (*F*(1,41)=40.157,*p*<.001; short vs long-term; *t*(52.05)=3.478,*p*=.001). The Third Thumb representation therefore became more disassociated from the controller toe movement following training. There was no difference in the distance between the Third Thumb and passive sensory feedback between groups (*F*(1,41)=1.528,*p*=.223).

These results, again echo the main findings from M1: following long-term training, the Third Thumb became more dissociated from its toe controller, without a corresponding change in its relationship to sensory feedback or the biological thumb.

#### Pattern Component Modelling

We repeated the same combined, independent and ceiling models structured in the main manuscript.

The combined model performed better than the independent model for explaining the Third Thumb representation for the short-term group (*p*<.001), while indicative evidence for an advantage following long-term experience was inconclusive (*p*=.056,BF_10_=1.389). With this borderline result, we then assessed the ability of the combined model to explain the variance in the data by comparing its fit to the noise ceiling. It was not significantly different for the short-term group (*p*=.138,BF_10_=.661) but the combined model fit was significantly below the noise ceiling for the long-term group (*p*=.009). To ensure we were choosing the best model to explain the data, we then compared to the independent model to the noise ceiling for the long-term group, however it was further below the noise ceiling than the combined (*p*<.001). When adding D1 as a biological component to the combined model, the two-component model performed better (*p*<.001). Therefore, we continued to explore the combined model that explained the Third Thumb activity just using its toe controller and sensory feedback components.

For reference, a biological body combined model was structured that explained the coordinated movements of D1 and Toe with the components of D1&Toe. When comparing this model against the Third Thumb model across groups (and accounting for their respective noise ceilings), there was a significant interaction between group and condition (*F*(1,126)=63.345,*p*<.001). Follow-up contrasts demonstrated that both the short- and long-term groups had a significant difference in the biological and Third Thumb models (both *p*<.001). However, in the short-term group the Third Thumb model performed better (Third Thumb estimated marginal mean (EMM)=-3174, D1 EMM=-3648) whilst in the long-term group, the D1 model performed better (Third Thumb EMM=-2962, D1 EMM=-2565).

To then understand how the constituent components combine to explain the Third Thumb representation, the relative weightings of each component in the model were assessed. When comparing groups, we found a significant interaction (Group X Component; *F*(1,41)=337.1,*p*<.001). Following short-term experience, the toe movement was the dominant component explaining the Third Thumb representation (significantly higher weighting; *t*(21)=10.15,*p*<.001). However, following longitudinal experience, this pattern was reversed (*t*(20)=16.66,*p*<.001), resulting in a significant increase in the weighting of the sensory feedback component (*t*(41)=18.36,*p*<.001), and a corresponding significant decrease for the weighting of the toe movement (*t*(41)=18.36,*p*<.001). This implies increased utilisation of Third Thumb sensory feedback on the hand by the S1 hand area.

Overall, PCM revealed the same experience-dependent re-weighting observed in M1, shifting from a predominantly toe-based contribution following short-term experience to a predominantly sensory-feedback contribution following long-term experience. However, unlike M1, the combined model remained significantly below the noise ceiling in the long-term S1 group, suggesting that the two constituent components provided a less complete account of Third Thumb-related representational structure in S1. The biological-body comparison also differed: whereas the difference between biological and Third Thumb models diminished with experience in M1, the models remained significantly different in S1, with their relative performance reversing across groups.

#### M1 Foot Area

Activity was significantly positive in the M1 foot area for all conditions involving toe movement across groups (Short-term: Toe movement (*t*(21)=9.169,*p*<.001), Third Thumb movement (*t*(21)=10.35,*p*<.001). Long-term: Toe movement (*t*(21)=8.915,*p*<.001), Third Thumb movement (*t*(21)=11.52,*p*<.001)), while the passive sensory feedback did not significantly activate the motor foot area (Short-term: (*t*(21)=.047,*p*=.963,BF_10_=.223). Long-term: (*t*(21)=.750,*p*=.462,BF_10_=.287)).

Importantly, there were no significant group differences in the activation of the foot area. There was a non-significant interaction when comparing the Toe and Third Thumb activity for both groups (Group X Condition: *F*(1,42)=2.404,*p*=.129). Follow-up analyses confirmed no group differences in activity for the toe movement (*t*(42)=.242,*p*=.810,BF_10_=.305) nor Third Thumb movement (*t*(42)=1.127,*p*=.266,BF_10_=.495).

#### Premotor Cortices

Given the role of premotor cortex in action selection, integration and preparation, we explored the univariate activity to see if there were any signs of experience-dependent differences. For group comparisons we continued to explore interaction terms relative to D1.

#### Premotor Ventral (PMv)

There was no significant difference between groups in the activity elicited during D1 movement (*t*(42)=1.264,*p*=.213,BF_10_=.564), and all conditions across groups elicited activity significantly above zero across the brain conditions of interest (D1, Third Thumb, sensory feedback and toe movement; all *p*<.001).

When comparing the experience groups, we found no significant difference in activity produced by toe movement in PMv (Group X Condition: *F*(1,42)=3.286,*p*=.077). However, we did find a significant interaction in the sensory feedback condition (*F*(1,42)=5.509,*p*=.024), but this was not driven by between-groups effects (feedback short vs long, *p*=.447). Finally, we also saw no differences for the Third Thumb activity (*F*(1,42)=.020,*p*=.889).

#### Premotor Dorsal (PMd)

There was no significant difference between groups in the activity elicited during D1 movement (*t*(42)=1.088,*p*=.283,BF_10_=.478), and all conditions across groups elicited activity significantly above zero across the brain conditions of interest (D1, Third Thumb, sensory feedback and toe movement; all *p* .002).

Within PMd, while we found no significant difference for the toe movement condition (Group X Condition: *F*(1,42)=3.727,*p*=.060) we did explore the trending p-value, and found it was not driven by any between-group effects (toe short vs long-term *p*=.427). We also found no differences for the sensory feedback (*F*(1,42)=.065,*p*=.799) nor Third Thumb (*F*(1,42)=.416,*p*=.523) conditions.

With no evidence for group differences, and no a priori hypothesis regarding premotor cortex, no further analyses were run.

### M1 Hand Area Replication with FIX

As an additional sensitivity analysis due to concerns about head movements, the long-term training dataset was denoised using MELODIC and FSL FIX with a study-specific classifier to assess whether movement-related artefacts could account for the observed effects. Applying this additional denoising did not alter the principal univariate, RSA or PCM results. All analyses reported in the main manuscript therefore use data processed without FIX, below is the replication with FIX.

#### Univariate

Moving D1 produced the expected activation in the M1 hand area (*t*(19)=10.81,*p*<.001), which was not significantly different from D1 activity found for the short-term training group (*t*(40)=1.213,*p*=.232,BF_10_=.508). However, with the Bayes factor only implying anecdotal evidence for similar activation amounts across groups, all further group comparisons explored interaction terms relative to D1.

Similarly to the short-term experience group, there was no significant activity in the motor hand area for toe movements (*t*(17)=.531,*p*=.602,BF_10_=.276), with no significant group differences (*F*(1,40)=1.249,*p*=.270,BF_10_=.211). Although the activity when simply receiving the sensory feedback from passive Thumb movement was still not above zero (*p*=.085,BF_10_=.980), we did, however, find significantly higher activation following long-term experience (Group X Condition interaction *F*(1,40)=11.31,*p*=.002; short-vs long-term training, *p*=.015). Importantly, Third Thumb movement in the long-term training group resulted in positive activity in the hand motor area *t*(17)=2.864,*p*=.011, resulting in significant group differences (*F*(1,40)=23.292,*p*<.001; short-vs long-term training, *p*<.001).

#### RSA

Between groups, the distance between D1 and the Third Thumb was not significantly different (*t*(38)=.787,*p*=.436,BF_10_=.396). Nevertheless, given the anecdotal Bayes factor, we assessed all further distances in relation to the Third Thumb to D1 distance.

We next assessed the distances between the Third Thumb and its two constituent components - toe movement and sensory feedback on the hand. The distance between the Third Thumb and toe movement was significantly greater following long-term experience compared to short (*F*(1,38)=29.191,*p*<.001; short vs long-term *t*(38)=4.280,*p*<.001). This increased separation from the toe condition was not accompanied by greater similarity to the sensory feedback (*F*(1,38)=.972,*p*=.330; short vs long-term *t(*60.42)=.069,*p*=.945,BF_10_=.311).

#### PCM

When assessing model performance, the combined model performed better than the independent model for the long-term (*p*=.035) experience group. The combined model was not significantly different from the noise ceiling (*p*=.449,BF_10_=.328). These model comparisons demonstrate the combined model sufficiently explains the Third Thumb representation and outperforms the alternative model that the representation is independent of its components.

We next assessed the relative weightings of each component in the model. Following longitudinal experience, we find a striking re-weighting of the components where the sensory feedback became the dominant component (*t*(17)=14.31,*p*<.001). Compared to short-term, we see a significant increase in the weighting of the sensory feedback component (*U*(38)=396,*p*<.001), and a corresponding significant decrease for the weighting of the toe movement (*U*(38)=0,*p*<.001).

When comparing the Third Thumb and biological combined models (and accounting for their respective noise ceilings) we found a significant interaction between group and condition (*F*(1,114)=40.473,*p*<.001). Follow-up contrasts demonstrated that this was driven by a difference in the biological and Third Thumb models for the short-term group (*p*<.001) not significant for the long-term experience group (*p*=.093, BF_10_≈.408).

Finally, we performed a follow-up analysis adding D1 to the combined Third Thumb model with the motor and sensory components to assess if adding this biological hand component improves the model fit. However, the overall model performance was best with just the two components constituting Third Thumb movement (*p*<.001 for both groups).

### Cortical Parcellation Analysis

In addition to the widespread effects observed for the Third Thumb-toe distance, significant experience-dependent differences were identified for the other representational distances in isolated cortical parcels. Following long-term training, the representational distance between the Third Thumb and biological thumb was significantly reduced in left s32, a region of the anterior cingulate cortex. The distance between the Third Thumb and sensory feedback was also significantly reduced in right V3B, a higher-order visual area within the dorsal visual stream. In both cases, this reflected greater representational similarity following long-term training. No other parcels survived comparison correction.

**Figure S5:**
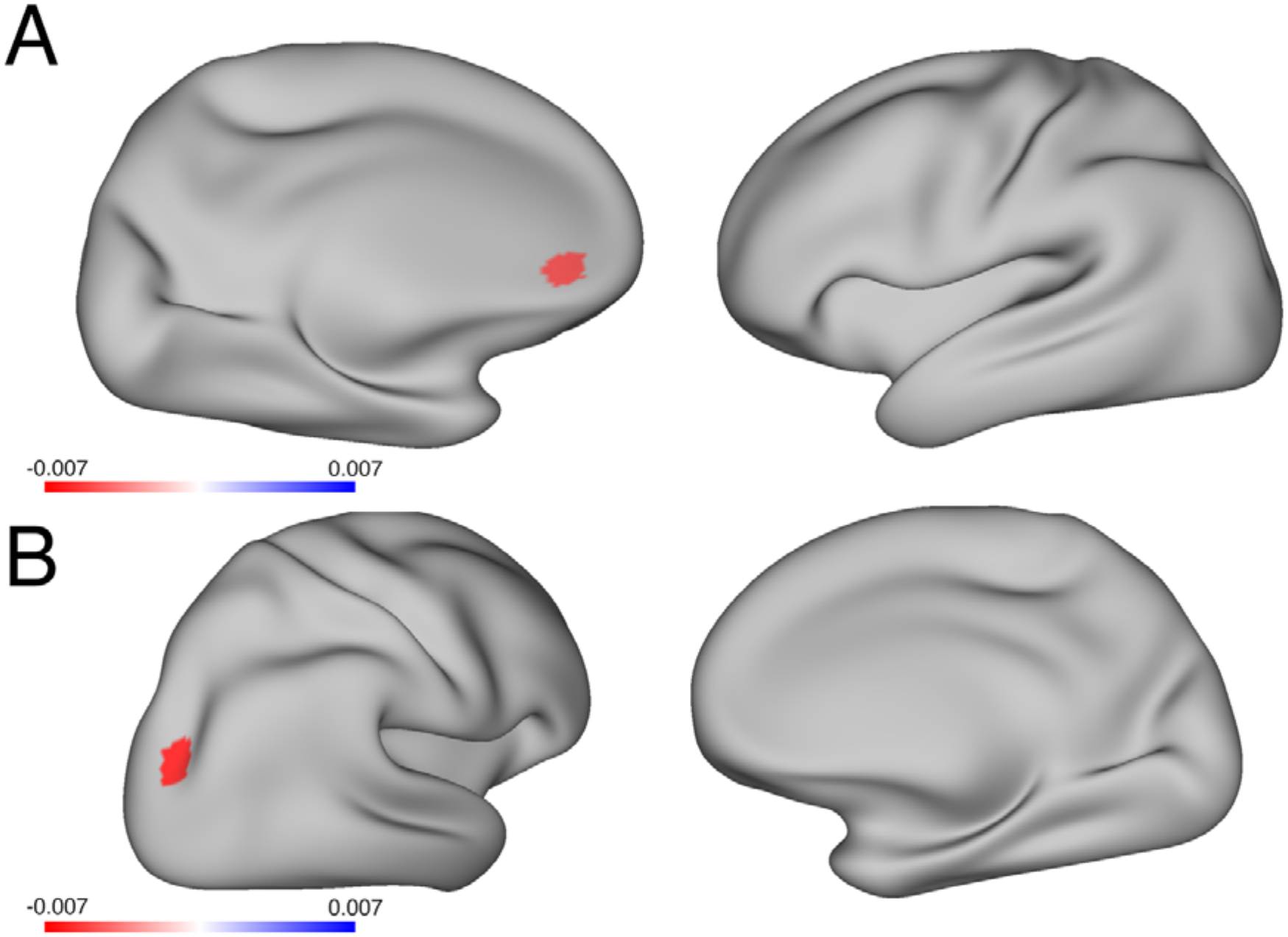
Experience-dependent differences in (A) Third Thumb-Bio Thumb and (B) Third Thumb-sensory feedback representational distance across the cortex. Cortical parcellations showing a significantly greater crossnobis distance between the Third Thumb and the respective condition in the short-term compared with the long-term training group following permutation-based maximum-|t| FWER correction across parcels (p<.05). Surface colour indicates the mean difference in crossnobis distance between groups (long-term training – short-term training), with larger values reflecting greater representational distance following long-term training

## Extended Methods

### Short-Term Training Tasks

Short-Term training participants were familiarised with the Third Thumb in the lab for approximately 90 minutes. Here, they started the session by training on a series of tasks designed to provide basic motor control of the device. These tasks were designed to be purposefully challenging when performed with only one hand. The task execution was restricted to the augmented (right) hand.

### Collaboration Task

The collaboration task involved using the Third Thumb in collaboration with a biological finger to pick up objects. A mixed jumble of wooden blocks was placed on the table. Participants were instructed to pick-up two blocks at a time, using the Third Thumb to support one of the blocks with another finger, to then build a 2×2 tower. The experimenter recorded the number of two-block floors built in one minute. The task was repeated five times.

### Individuation Task

The individuation task worked on fine motor control with the Third Thumb, while having the hand fully occupied with a task-irrelevant object. Participants were given rolls of tape, a foam ball and a wooden pole fixed on the desk. While holding the ball with their biological fingers, participants had to use active movements with the Third Thumb to pick up the tape rolls and place them on the wooden pole. The experimenter recorded how many tapes could be placed on the wooden pole in one minute. The task was repeated five times.

### Coordination Task

The coordination task was a finger opposition task. Participants were seated in front of a computer screen that displayed task stimuli. They were instructed to move the Third Thumb to touch the tip of a randomly specified finger on the augmented (right) hand; emphasis was placed on a specific tip-to-tip movement. Participants were instructed to make as many successful hits within a one-minute block. Participants completed a total of 10 blocks. A MATLAB script was used to randomly select a target finger (‘thumb’,’index’,’middle’,’ring’ or ‘little’) and to display the finger name on the computer in front of the participant. The experimenter manually advanced the program to the next stimulus when participants successfully touched the tip of the target finger with the Third Thumb.

### Free Form Tasks

Following the structured training tasks, participants were given an array of objects (an alphabet puzzle board, a piggy bank and coins, a zipped bag, a peg board, and a children’s book). They had approximately 20 minutes to explore and use the Third Thumb however they want. They were encouraged to explore and challenge themselves, but they were free to do whichever tasks they preferred, as these were non-scored training tasks.

## References

Arbuckle, S. A., Yokoi, A., Pruszynski, J. A., & Diedrichsen, J. (2019). Stability of representational geometry across a wide range of fMRI activity levels. NeuroImage, 186, 155–163. 10.1016/j.neuroimage.2018.11.002

Bassett, D. S., Yang, M., Wymbs, N. F., & Grafton, S. T. (2015). Learning-induced autonomy of sensorimotor systems. Nature Neuroscience, 18(5), 744–751. 10.1038/NN.3993

Berlot, E., Popp, N. J., & Diedrichsen, J. (2020). A critical re-evaluation of fMRI signatures of motor sequence learning. ELife, 9, 1–24. 10.7554/ELIFE.55241

Clode, D., Dowdall, L., da Silva, E., Selén, K., Cowie, D., Dominijanni, G., & Makin, T. R. (2024). Evaluating initial usability of a hand augmentation device across a large and diverse sample. Science Robotics, 9(90). 10.1126/scirobotics.adk5183

Dall’Orso, S., Steinweg, J., Allievi, A. G., Edwards, A. D., Burdet, E., & Arichi, T. (2018). Somatotopic Mapping of the Developing Sensorimotor Cortex in the Preterm Human Brain. Cerebral Cortex (New York, N.Y. : 1991), 28(7), 2507–2515. 10.1093/CERCOR/BHY050

Dash, D., Iwane, F., Hayward, W., Salamanca-Giron, R. F., Bönstrup, M., Buch, E. R., & Cohen, L. G. (2025). Sequence action representations contextualize during early skill learning. ELife, 13, RP102475. 10.7554/ELIFE.102475

Dempsey-Jones, H., Harrar, V., Oliver, J., Johansen-Berg, H., Spence, C., & Makin, T. R. (2016). Transfer of tactile perceptual learning to untrained neighboring fingers reflects natural use relationships. Journal of Neurophysiology, 115(3), 1088–1097. 10.1152/JN.00181.2015

Deo, D. R., Okorokova, E. V., Pritchard, A. L., Hahn, N. V., Card, N. S., Nason-Tomaszewski, S. R., Jude, J., Hosman, T., Choi, E. Y., Qiu, D., Meng, Y., Wairagkar, M., Nicolas, C., Kamdar, F. B., Iacobacci, C., Acosta, A., Hochberg, L. R., Cash, S. S., Williams, Z. M., … Willett, F. R. (2026). A mosaic of whole-body representations on the human precentral gyrus. Nature 2026, 1–8. 10.1038/s41586-026-10653-x

Diedrichsen, J., Ridgway, G. R., Friston, K. J., & Wiestler, T. (2011). Comparing the similarity and spatial structure of neural representations: A pattern-component model. NeuroImage, 55(4), 1665–1678. 10.1016/J.NEUROIMAGE.2011.01.044

Diedrichsen, J., Yokoi, A., & Arbuckle, S. A. (2018). Pattern component modeling: A flexible approach for understanding the representational structure of brain activity patterns. NeuroImage, 180(Pt A), 119–133. 10.1016/j.neuroimage.2017.08.051

Dominijanni, G., Shokur, S., Salvietti, G., Buehler, S., Palmerini, E., Rossi, S., De Vignemont, F., d’Avella, A., Makin, T. R., Prattichizzo, D., & Micera, S. (2021). The neural resource allocation problem when enhancing human bodies with extra robotic limbs. Nature Machine Intelligence 2021 3:10, 3(10), 850–860. 10.1038/s42256-021-00398-9

Dowdall, L., Molina-Sanchez, M., Dominijanni, G., Silva, E. da, Pavalkyte, V., Jugovic, E., Bianchi, M., Iida, F., Clode, D., & Makin, T. R. (2025). Developing a Sensory Representation of an Artificial Body Part. BioRxiv, 2025.06.16.658246. 10.1101/2025.06.16.658246

Eden, J., Bräcklein, M., Ibáñez, J., Barsakcioglu, D. Y., Di Pino, G., Farina, D., Burdet, E., & Mehring, C. (2022). Principles of human movement augmentation and the challenges in making it a reality. Nature Communications, 13(1), 1–13. 10.1038/S41467-022-28725-7

Ejaz, N., Hamada, M., & Diedrichsen, J. (2015). Hand use predicts the structure of representations in sensorimotor cortex. Nature Neuroscience, 18(7), 1034– 1040. 10.1038/nn.4038

Fischl, B. (2012). FreeSurfer. NeuroImage, 62(2), 774–781. 10.1016/J.NEUROIMAGE.2012.01.021

Fischl, B., Rajendran, N., Busa, E., Augustinack, J., Hinds, O., Yeo, B. T. T., Mohlberg, H., Amunts, K., & Zilles, K. (2008). Cortical folding patterns and predicting cytoarchitecture. Cerebral Cortex (New York, N.Y. : 1991), 18(8), 1973–1980. 10.1093/CERCOR/BHM225

Hahamy, A., Macdonald, S. N., van den Heiligenberg, F., Kieliba, P., Emir, U., Malach, R., Johansen-Berg, H., Brugger, P., Culham, J. C., & Makin, T. R. (2017). Representation of Multiple Body Parts in the Missing-Hand Territory of Congenital One-Handers. Current Biology, 27(9), 1350–1355. 10.1016/j.cub.2017.03.053

Howell, P., Rabe, F., Meneghin, C., Schading-Sassenhausen, S., Schone, H., Sonar, H. A., Paik, J., Freund, P., Wenderoth, N., & Kikkert, S. (2026). Revisiting Face-to-Hand Area Remapping in the Human Primary Somatosensory Cortex after a Cervical Spinal Cord Injury. The Journal of Neuroscience : The Official Journal of the Society for Neuroscience, 46(32), e0644252026. 10.1523/JNEUROSCI.0644-25.2026

Huber, L., Finn, E. S., Handwerker, D. A., Bönstrup, M., Glen, D. R., Kashyap, S., Ivanov, D., Petridou, N., Marrett, S., Goense, J., Poser, B. A., & Bandettini, P. A. (2020). Sub-millimeter fMRI reveals multiple topographical digit representations that form action maps in human motor cortex. NeuroImage, 208. 10.1016/j.neuroimage.2019.116463

Hussain, I., Santarnecchi, E., Leo, A., Ricciardi, E., Rossi, S., & Prattichizzo, D. (2017). A magnetic compatible supernumerary robotic finger for functional magnetic resonance imaging (fMRI) acquisitions: Device description and preliminary results. IEEE … International Conference on Rehabilitation Robotics : [Proceedings], 2017, 1177–1182. 10.1109/ICORR.2017.8009409

Jenkinson, M., Bannister, P., Brady, M., & Smith, S. (2002). Improved optimization for the robust and accurate linear registration and motion correction of brain images. NeuroImage, 17(2), 825–841. 10.1016/S1053-8119(02)91132-8

Jenkinson, M., Beckmann, C. F., Behrens, T. E. J., Woolrich, M. W., & Smith, S. M. (2012). FSL. NeuroImage, 62(2), 782–790. 10.1016/J.NEUROIMAGE.2011.09.015

Jenkinson, M., & Smith, S. (2001). A global optimisation method for robust affine registration of brain images. Medical Image Analysis, 5(2), 143–156. 10.1016/S1361-8415(01)00036-6

Kieliba, P., Clode, D., Maimon-Mor, R. O., & Makin, T. R. (2021). Robotic hand augmentation drives changes in neural body representation. Science Robotics, 6(54). 10.1126/SCIROBOTICS.ABD7935

Kikkert, S., Kolasinski, J., Jbabdi, S., Tracey, I., Beckmann, C. F., Berg, H. J., & Makin, T. R. (2016). Revealing the neural fingerprints of a missing hand. ELife, 5(AUGUST). 10.7554/ELIFE.15292

Kikkert, S., Pfyffer, D., Verling, M., Freund, P., & Wenderoth, N. (2021). Finger somatotopy is preserved after tetraplegia but deteriorates over time. ELife, 10. 10.7554/ELIFE.67713

Kleinschmidt, A., Nitschke, M., & Frahm, J. (1997). Somatotopy in the human motor cortex hand area. A high-resolution functional MRI study. The European Journal of Neuroscience, 9(10), 2178–2186. 10.1111/J.1460-9568.1997.TB01384.X

Kolasinski, J., Makin, T. R., Logan, J. P., Jbabdi, S., Clare, S., Stagg, C. J., & Johansen-Berg, H. (2016). Perceptually relevant remapping of human somatotopy in 24 hours. ELife, 5(DECEMBER2016). 10.7554/ELIFE.17280

Kolasinski, X., Makin, T. R., Jbabdi, S., Clare, S., Stagg, C. J., & Johansen-Berg, H. (2016). Investigating the Stability of Fine-Grain Digit Somatotopy in Individual Human Participants. The Journal of Neuroscience : The Official Journal of the Society for Neuroscience, 36(4), 1113–1127. 10.1523/JNEUROSCI.1742-15.2016

Krakauer, J. W., & Carmichael, S. T. (2017). Broken Movement: The Neurobiology of Motor Recovery after Stroke. Broken Movement. 10.7551/MITPRESS/9310.001.0001

Kriegeskorte, N., Mur, M., & Bandettini, P. (2008). Representational similarity analysis - connecting the branches of systems neuroscience. Frontiers in Systems Neuroscience, 2(NOV), 249. 10.3389/NEURO.06.004.2008

Liu, Y., Huang, S., Wang, Z., Ji, F., & Ming, D. (2022). Functional Reorganization After Four-Week Brain-Computer Interface-Controlled Supernumerary Robotic Finger Training: A Pilot Study of Longitudinal Resting-State fMRI. Frontiers in Neuroscience, 15. 10.3389/FNINS.2021.766648

Maimon-Mor, R. O., Schone, H. R., Slater, D. H., Faisal, A. A., & Makin, T. R. (2021). Early life experience sets hard limits on motor learning as evidenced from artificial arm use. ELife, 10. 10.7554/ELIFE.66320

Makin, T. R., Cramer, A. O., Scholz, J., Hahamy, A., Slater, D. H., Tracey, I., & Johansen-Berg, H. (2013). Deprivation-related and use-dependent plasticity go hand in hand. ELife, 2(November2013), 1–15. 10.7554/ELIFE.01273

Mancini, F., Wang, A. P., Schira, M. M., Isherwood, Z. J., McAuley, J. H., Iannetti, G. D., Sereno, M. I., Moseley, G. L., & Rae, C. D. (2019). Fine-Grained Mapping of Cortical Somatotopies in Chronic Complex Regional Pain Syndrome. Journal of Neuroscience, 39(46), 9185–9196. 10.1523/JNEUROSCI.2005-18.2019

Meltzoff, A. N., Saby, J. N., & Marshall, P. J. (2019). Neural representations of the body in 60-day-old human infants. Developmental Science, 22(1). 10.1111/DESC.12698

Molina-Sanchez, M., Dowdall, L., Dominijanni, G., Pavalkyte, V., Gallay, C., da Silva, E., Clode, D., & Makin, T. R. (2026). Broad skill generalization with an extra robotic body part. Current Biology, 36(5), 1339–1346.e4. 10.1016/j.cub.2026.01.069

Muret, D., Root, V., Kieliba, P., Clode, D., & Makin, T. R. (2022). Beyond body maps: Information content of specific body parts is distributed across the somatosensory homunculus. Cell Reports, 38(11). 10.1016/j.celrep.2022.110523

Nili, H., Wingfield, C., Walther, A., Su, L., Marslen-Wilson, W., & Kriegeskorte, N. (2014). A toolbox for representational similarity analysis. PLoS Computational Biology, 10(4). 10.1371/JOURNAL.PCBI.1003553

Ogawa, K., Mitsui, K., Imai, F., & Nishida, S. (2019). Long-term training-dependent representation of individual finger movements in the primary motor cortex. NeuroImage, 202, 116051. 10.1016/J.NEUROIMAGE.2019.116051

Omrani, M., Murnaghan, C. D., Pruszynski, J. A., & Scott, S. H. (2016). Distributed task-specific processing of somatosensory feedback for voluntary motor control. ELife, 5(APRIL2016). 10.7554/ELIFE.13141

Penfield, W., & Boldrey, E. (1937). SOMATIC MOTOR AND SENSORY REPRESENTATION IN THE CEREBRAL CORTEX OF MAN AS STUDIED BY ELECTRICAL STIMULATION. Brain, 60(4), 389–443. 10.1093/BRAIN/60.4.389

Power, J. D., Mitra, A., Laumann, T. O., Snyder, A. Z., Schlaggar, B. L., & Petersen, S. E. (2013). Methods to detect, characterize, and remove motion artifact in resting state fMRI. NeuroImage, 84, 10.1016/j.neuroimage.2013.08.048. 10.1016/J.NEUROIMAGE.2013.08.048

Prattichizzo, D., Pozzi, M., Baldi, T. L., Malvezzi, M., Hussain, I., Rossi, S., & Salvietti, G. (2021). Human augmentation by wearable supernumerary robotic limbs: review and perspectives. Progress in Biomedical Engineering, 3(4), 042005. 10.1088/2516-1091/AC2294

Pruszynski, J. A., Omrani, M., & Scott, S. H. (2014). Goal-dependent modulation of fast feedback responses in primary motor cortex. The Journal of Neuroscience : The Official Journal of the Society for Neuroscience, 34(13), 4608–4617. 10.1523/JNEUROSCI.4520-13.2014

Raffin, E., & Siebner, H. R. (2019). Use-Dependent Plasticity in Human Primary Motor Hand Area: Synergistic Interplay Between Training and Immobilization. Cerebral Cortex (New York, N.Y. : 1991), 29(1), 356–371. 10.1093/CERCOR/BHY226

Rao, S. M., Binder, J. R., Hammeke, T. A., Bandettini, P. A., Bobholz, J. A., Frost, J. A., Myklebust, B. M., Jacobson, R. D., & Hyde, J. S. (1995). Somatotopic mapping of the human primary motor cortex with functional magnetic resonance imaging. Neurology, 45(5), 919–924. 10.1212/WNL.45.5.919

Rossi, S., Salvietti, G., Neri, F., Romanella, S. M., Cinti, A., Sinigaglia, C., Ulivelli, M., Lisini Baldi, T., Santarnecchi, E., & Prattichizzo, D. (2021). Emerging of new bioartificial corticospinal motor synergies using a robotic additional thumb. Scientific Reports 2021 11:1, 11(1), 18487-. 10.1038/s41598-021-97876-2

Schone, H. R., Maimon-Mor, R. O., Kollamkulam, M., Szymanska, M. A., Gerrand, C., Woollard, A., Kang, N. V., Baker, C. I., & Makin, T. R. (2025). Stable cortical body maps before and after arm amputation. Nature Neuroscience 2025 28:10, 28(10), 2015–2021. 10.1038/s41593-025-02037-7

Shelchkova, N. D., Downey, J. E., Greenspon, C. M., Okorokova, E. V., Sobinov, A. R., Verbaarschot, C., He, Q., Sponheim, C., Tortolani, A. F., Moore, D. D., Kaufman, M. T., Lee, R. C., Satzer, D., Gonzalez-Martinez, J., Warnke, P. C., Miller, L. E., Boninger, M. L., Gaunt, R. A., Collinger, J. L., … Bensmaia, S. J. (2023). Microstimulation of human somatosensory cortex evokes task-dependent, spatially patterned responses in motor cortex. Nature Communications 2023 14:1, 14(1), 7270-. 10.1038/s41467-023-43140-2

Smith, S. M. (2002). Fast robust automated brain extraction. Human Brain Mapping, 17(3), 143–155. 10.1002/HBM.10062

Tucciarelli, R., Bird, L., Straka, Z., Szymanska, M., Kollamkulam, M., Sonar, H. A., Paik, J., Clode, D., Hoffmann, M., Cowie, D., & Makin, T. R. (2026). Global remapping of the sensory homunculus emerges early in childhood development. Nature Communications 2025 17:1, 17(1), 1591-. 10.1038/s41467-025-66539-5

Wesselink, D. B., Heiligenberg, F. M. Van Den, Ejaz, N., Dempsey-Jones, H., Cardinali, L., Tarall-Jozwiak, A., Diedrichsen, J., & Makin, T. R. (2019). Obtaining and maintaining cortical hand representation as evidenced from acquired and congenital handlessness. ELife, 8. 10.7554/ELIFE.37227

Wesselink, D. B., Sanders, Z. B., Edmondson, L. R., Dempsey-Jones, H., Kieliba, P., Kikkert, S., Themistocleous, A. C., Emir, U., Diedrichsen, J., Saal, H. P., & Makin, T. R. (2022). Malleability of the cortical hand map following a finger nerve block. Science Advances, 8(16), 2393. 10.1126/SCIADV.ABK2393

Wesselink, D., & Maimon-Mor, R. (2017). *rsatoolbox, version* 20fbe05.github.

Wetzels, R., Matzke, D., Lee, M. D., Rouder, J. N., Iverson, G. J., & Wagenmakers, E. J. (2011). Statistical Evidence in Experimental Psychology: An Empirical Comparison Using 855 t Tests. Perspectives on Psychological Science : A Journal of the Association for Psychological Science, 6(3), 291–298. 10.1177/1745691611406923

Wiestler, T., & Diedrichsen, J. (2013). Skill learning strengthens cortical representations of motor sequences. ELife, 2(2). 10.7554/ELIFE.00801

Wiestler, T., Waters-Metenier, S., & Diedrichsen, J. (2014). Effector-independent motor sequence representations exist in extrinsic and intrinsic reference frames. The Journal of Neuroscience : The Official Journal of the Society for Neuroscience, 34(14), 5054–5064. 10.1523/JNEUROSCI.5363-13.2014

Wu, F., & Asada, H. (2014). Supernumerary Robotic Fingers: An Alternative Upper-Limb Prosthesis. ASME 2014 Dynamic Systems and Control Conference, DSCC 2014, 2. 10.1115/DSCC2014-6017

Yokoi, A., Arbuckle, S. A., & Diedrichsen, J. (2018). The Role of Human Primary Motor Cortex in the Production of Skilled Finger Sequences. The Journal of Neuroscience : The Official Journal of the Society for Neuroscience, 38(6), 1430–1442. 10.1523/JNEUROSCI.2798-17.2017

Yousry, T. A., Schmid, U. D., Alkadhi, H., Schmidt, D., Peraud, A., Buettner, A., & Winkler, P. (1997). Localization of the motor hand area to a knob on the precentral gyrus. A new landmark. Brain : A Journal of Neurology, 120 ( Pt 1)(1), 141–157. 10.1093/BRAIN/120.1.141

